# Trans-branching of polyubiquitin chains orchestrates the DNA replication stress response

**DOI:** 10.64898/2026.09.01.748521

**Authors:** Kirill Petriukov, Christian Renz, Markus S. Schraft, Ivan Mikicic, Nils C. Krapoth, Abhik Thapa, Petra Beli, Helle D. Ulrich

## Abstract

Polyubiquitin chain geometry dictates functional consequences of ubiquitylation (*1, 2*). Although branched polyubiquitin chains are abundant in cells, little is known about their functions (*3–5*). Here we show that branching on the DNA replication factor PCNA, mediated by the ubiquitin-conjugating enzyme UBE2K and involving lysines 63 and 48 of ubiquitin, orchestrates the sequence of events in response to replication stress. By inducing VCP-dependent extraction of PCNA from chromatin, branching promotes re-priming of stalled forks and necessitates a BRCA1-dependent pathway of daughter-strand gap repair. Our study identifies hyper-accumulation of daughter-strand gaps as the mechanistic basis underlying the toxicity of inhibitors of the PCNA-specific isopeptidase, USP1, in BRCA1-deficient cells. Moreover, an unexpected preference of UBE2K to operate *in trans* suggests a general timing mechanism to organize hierarchies amongst ubiquitin signals.

## Main

Polyubiquitin chains convey a variety of cellular signals, both degradative and non-degradative, via their distinct geometries (*1, 2*). These are determined by the enzymes required for their assembly, the ubiquitin-conjugating enzymes (E2s) and ubiquitin protein ligases (E3s), and may involve mixed linkages or even branching, when multiple lysine (K) residues engage in chain elongation. Our insight into the signaling functions of homotypic polyubiquitin chains is rapidly growing, but we are only beginning to understand the implications of branched chains, which account for as much as 10-20% of all chain-associated ubiquitin (*3–5*). In most reported cases, branching appears to promote proteasomal targeting by amplifying the ubiquitin signal (*6–9*), but it may also protect ubiquitin chains from attack by deubiquitylating enzymes (DUBs) (*10, 11*) or convey two independent non-proteolytic signals (*12*). A prominent effector of polyubiquitylation is VCP (p97), a ubiquitin-dependent segregase that cooperates with a large set of ubiquitin-binding adaptors and generally facilitates degradation via disassembly of protein complexes or their extraction from membranes (*13*). Depending on its adaptors, VCP may prefer either K48-linked chains or branched or complex ubiquitin conjugates (*6, 8, 14*).

The ubiquitin system contributes to many aspects of genome maintenance (*15*). Management of DNA replication stress is paramount to genome stability as it ensures efficient and accurate genome duplication in proliferating cells. In eukaryotes, the response to replication fork-stalling lesions is governed by mono-and polyubiquitylation of the essential replication factor, proliferating cell nuclear antigen (PCNA) (*16*). Monoubiquitylation of PCNA by the E3, RAD18, at an invariant lysine, K164, promotes translesion synthesis by damage-tolerant DNA polymerases (*17, 18*). PCNA K63-polyubiquitylation is much less well understood but has been implicated in fork reversal in human cells (*19, 20*). Moreover, PCNA modifications play a controversial role in relation to the breast cancer gene, BRCA1. In BRCA1-deficient cells, accumulation of ubiquitylated PCNA, imposed by inhibition or depletion of the PCNA-specific DUB, USP1, causes RAD18-dependent replication problems and loss of viability (*21, 22*). This synthetic lethal relationship has important clinical implications as USP1 inhibitors are being developed for therapeutic purposes (*23, 24*). Elucidating its mechanistic basis is critical to understand how the supportive function of PCNA ubiquitylation in genome maintenance can be reconciled with its detrimental effects in BRCA1-deficient cancers.

### UBE2K promotes K63-to-K48 trans-branching

Consistent with previous reports (*21, 22*), inhibition of USP1 (USP1i) in a BRCA1-deficient cancer cell line, MDA-MB-436, resulted in massive PCNA degradation, DNA damage signaling, and loss of viability in a RAD18-dependent manner (Fig. 1a, Extended data Fig. 1a,b). Depletion of the E3, RFWD3, previously shown to be required for PCNA polyubiquitylation (*25*), also suppressed PCNA degradation and checkpoint activation (Fig. 1a), arguing against an involvement of monoubiquitylation. Intriguingly, an unrelated, K48-selective E2, UBE2K, had been implicated in the toxic effects of USP1i (*22*); accordingly, its depletion suppressed the observed phenotypes in our hands (Fig. 1a, Extended Data Fig. 1b). Moreover, overexpression of UBE2K enhanced PCNA degradation, damage signaling and loss of viability, while a catalytically inactive mutant had the opposite effect (Extended Data Fig. 1c,d). We therefore asked whether UBE2K can directly ubiquitylate PCNA. *In vitro* assays with recombinant proteins demonstrated UBE2K activity only towards K63-di-or polyubiquitylated, but not unmodified or monoubiquitylated PCNA (Fig. 1b).

**Fig. 1.**
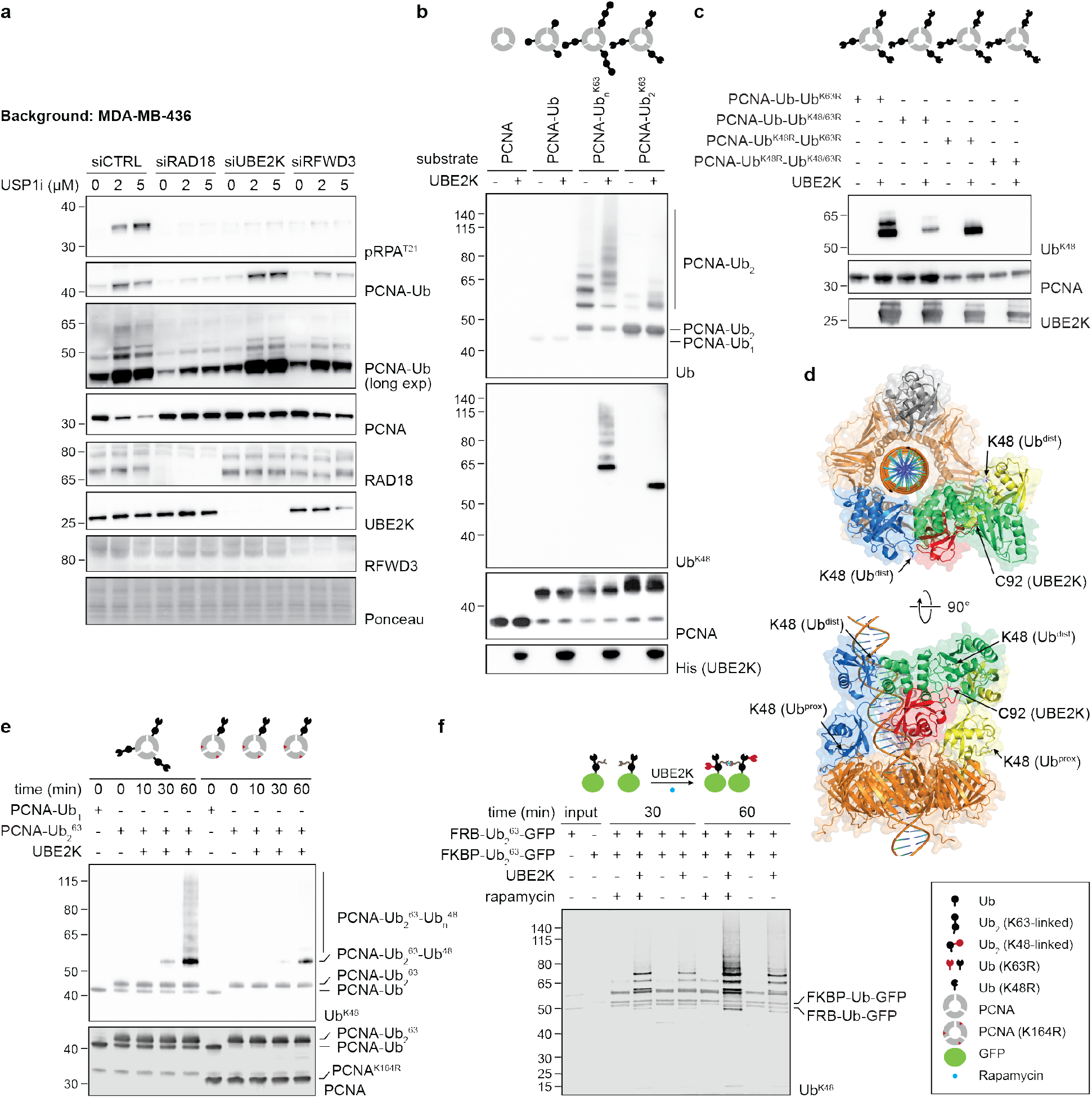
UBE2K promotes trans-branching. (**a**) Depletion of RAD18, UBE2K, or RFWD3 prevents checkpoint activation and PCNA degradation in BRCA1-deficient MDA-MB-436 cells upon USP1i (30 µM ML-323) for 72 h (pRPA^T21^: RPA32 phosphorylated at T21). (**b**) UBE2K affords K63-to-K48 branching of di-and polyubiquitylated PCNA *in vitro* (Ub^K48^: K48-linkage-specific anti-ubiquitin antibody). (**c**) UBE2K preferentially modifies the distal ubiquitin moieties of diubiquitylated PCNA. (**d**) Trans-branching of diubiquitylated, DNA-loaded PCNA by UBE2K is sterically plausible. Top and side views of an AlphaFold 3 model, suggesting a bridging of UBE2K between neighboring diubiquitin units on PCNA (UBE2K-bound Ub_2_: blue; acceptor Ub_2_: yellow; donor Ub: red). Note that only part of the DNA is shown. (**d**) At equal diubiquitin concentration, UBE2K prefers to act on PCNA trimers carrying diubiquitin chains on multiple subunits. (**f**) UBE2K preferentially branches K63-chains proximal to other chains. Branching was assessed *in vitro* on K63-diubiquitylated Ub-GFP constructs dimerizable by rapamycin.

UBE2K’s budding yeast homolog, Ubc1, modifies K48 of the proximal ubiquitin within an unanchored K63-linked ubiquitin dimer (*26*). Surprisingly, using fully K63-diubiquitylated PCNA as a substrate, we found that UBE2K modified K48 of both ubiquitin moieties, albeit with a strong preference for the distal unit (Fig. 1c, Extended Data Fig. 1e). Considering the trimeric nature of PCNA, we reasoned that the E2 might act *in trans*, modifying a diubiquitin chain on one subunit while binding to the chain present on another subunit. In this scenario, the distal ubiquitin might be more accessible to the E2’s catalytic site. An AlphaFold 3 (*27*) model of diubiquitylated, trimeric PCNA in complex with DNA and UBE2K, supported this arrangement (Fig. 1d, Extended Data Fig. 2). We therefore produced PCNA trimers carrying either a single or up to three diubiquitin chains (Extended Data Fig. 3a). At equal overall concentration of diubiquitin, UBE2K was considerably more active towards multiply modified trimers, indicating that – consistent with trans-branching – UBE2K’s activity is stimulated by additional K63-chains in its proximity (Fig. 1e).

When UBE2K was applied to K63-polyubiquitylated GFP fused to a pair of rapamycin-inducible dimerization domains (Extended Data Fig. 3b), branching via K48 was strongly stimulated by dimerization (Fig. 1f, Extended Data Fig. 3c), indicating that trans-branching is a substrate-independent property of this E2. Moreover, on asymmetric pairs of dimerizable mono-and K63-diubiquitylated GFP, UBE2K preferentially extended the monoubiquitylated species (Extended Data Fig. 3d), confirming the importance of non-covalent binding to the K63-chain. We conclude that UBE2K generally prefers to act *in trans* on substrates or complexes carrying multiple K63-chains.

### UBE2K-dependent branching induces VCP-mediated PCNA extraction from chromatin

To interrogate the fate and functional consequences of ubiquitylated PCNA in isolation from the general replication stress response, we selectively induced unscheduled PCNA K63-polyubiquitylation by means of a custom, PCNA-and linkage-specific E3, PIP-E3^63^ (*28*), in non-transformed RPE1 hTERT FlpIn cells (Fig. 2a, Extended Data Fig. 4a). Doxycycline (Dox)-induced expression of a stably integrated PIP-E3^63^ construct caused a severe and persistent cell cycle arrest in early S phase within 24 hours (Fig. 2b, Extended Data Fig. 4b). Mutant E3s lacking the PCNA interaction motif (ΔPIP), harboring a catalytically sub-active point mutation (I227A), or carrying a complete deletion of the RING domain (ΔRING), had no effect, indicating a requirement for both PCNA interaction and catalytic activity (Fig. 2b). DNA fiber assays revealed a massive reduction in replication fork speed (Fig. 2c), and phosphorylation of checkpoint kinases, CHK1 and CHK2, as well as RPA32, suggested an excessive exposure of single-stranded DNA (Fig. 2d). At later timepoints, a γH2AX signal indicated chromosome breakage. Again, PIP-E3^63^ mutants elicited no response (Fig. 2e). Moreover, expression of PIP-E3^63^ but not of its inactive mutants drastically interfered with cell viability (Extended Data Fig. 4c). In a PCNA(K164R) mutant cell line (*29*) or upon depletion of RAD18, PIP-E3^63^ did not induce cell cycle arrest or checkpoint activation (Extended Data Fig. 4d-h), thus excluding off-target effects. Thus, excessive PCNA K63-polyubiquitylation mimics the toxic effects of inhibiting PCNA deubiquitylation in BRCA1-deficient cells as described above.

**Fig. 2.**
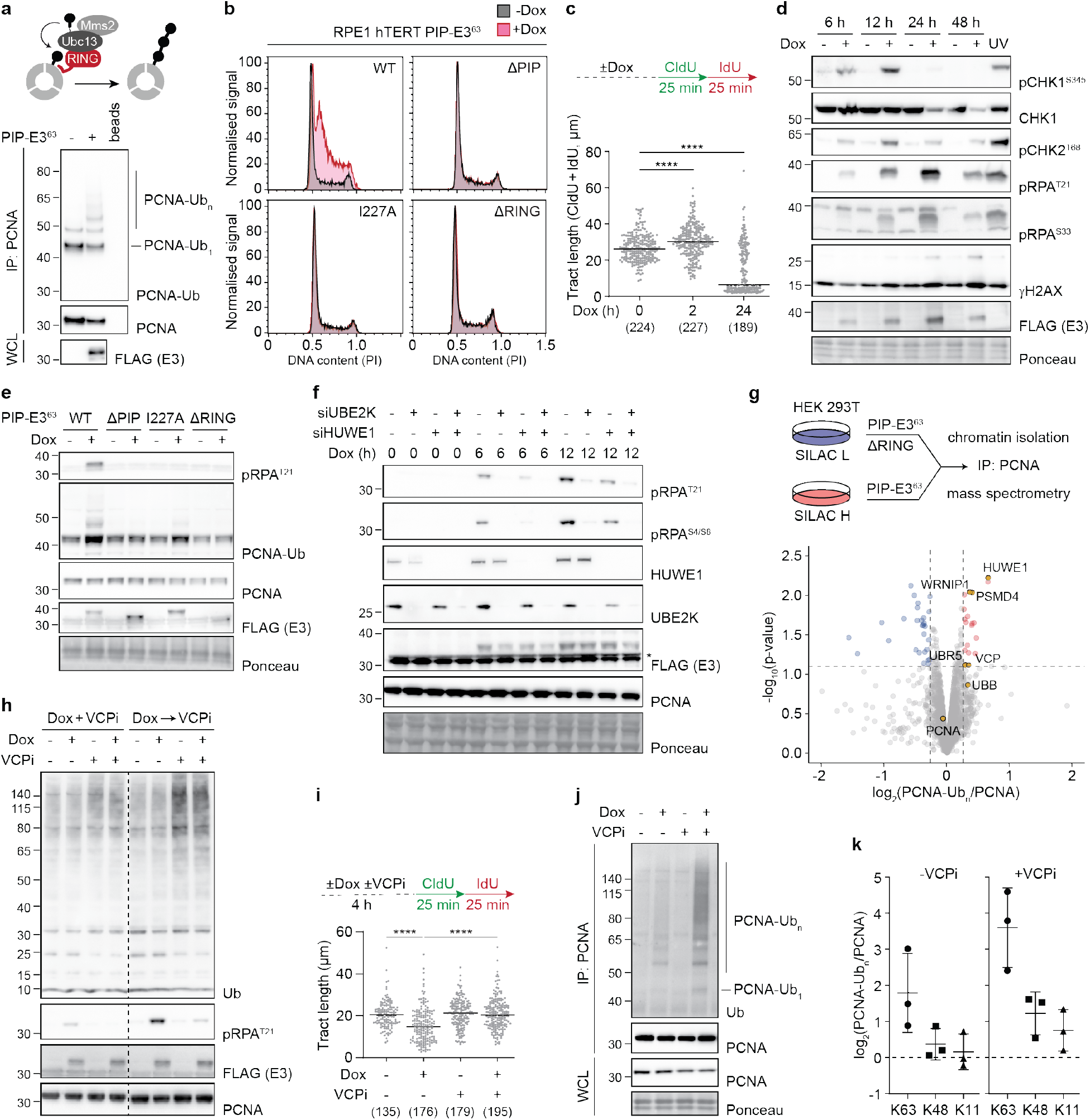
PCNA polyubiquitylation induces VCP-dependent extraction from chromatin. (**a**) Top: schematic view of PIP-E3^63^ (red) for induced K63-polyubiquitylation of PCNA. Bottom: PIP-E3^63^ affords PCNA polyubiquitylation after transient expression for 24 h in HEK 293T cells. Ubiquitylation of PCNA was analyzed with a PCNA-Ub-specific antibody after PCNA immunoprecipitation (IP) from the chromatin fraction, and expression of the E3 was analyzed in whole cell lysates (WCL) by western blotting. (**b**) PIP-E3^63^ causes a cell cycle arrest in early S phase. RPE1 hTERT FlpIn cells harboring stably integrated PIP-E3^63^ or indicated mutants were treated with 1 µg/ml doxycycline (Dox) 24 h prior to cell cycle analysis (PI: propidium iodide). (**c**) PIP-E3^63^ causes replication stalling. DNA fiber assays were performed on RPE1 hTERT PIP-E3^63^ FlpIn cells treated with 1 µg/ml Dox for 2 or 24 h. Only tracts containing green and red signals were analyzed. A representative experiment of two replicates is shown. Significance levels were calculated using the two-tailed Mann-Whitney test from the indicated number of fibers per sample (****: p<0.0001). (**d**) PIP-E3^63^ activates damage signaling. RPE1 hTERT PIP-E3^63^ FlpIn cells were treated with 1 µg/ml Dox for the indicated periods. Checkpoint markers and E3 expression were analyzed by western blotting of total cell extracts. An extract of cells treated with 40 J/m^2^ UV, followed by 4 h of recovery, served as positive control for checkpoint activation. (**e**) PIP-E3^63^-induced damage signaling requires E3 activity. Checkpoint activation and PCNA ubiquitylation were analyzed as in panel **d** (24 h Dox), using RPE1 hTERT FlpIn cells harboring wildtype (WT) or indicated E3 mutants. (**f**) UBE2K is required for PIP-E3^63^-induced checkpoint activation. HUWE1 and UBE2K were transiently depleted in RPE1 hTERT FlpIn PIP-E3^63^ cells. 72 h after transfection, cells were treated with 1 µg/ml Dox for the indicated periods. Checkpoint activation, depletion efficiency, and PIP-E3^63^ expression were analyzed by western blotting of total cell extracts. The asterisk marks a cross-reacting band. (**g**) SILAC-based mass spectrometry identifies interactors of chromatin-bound K63-polyubiquitylated PCNA. The volcano plot shows proteins depleted for interaction (blue) or preferentially interacting with polyubiquitylated PCNA (red). Selected interactors are highlighted and labeled (UBB: ubiquitin). (**h**) VCPi prevents PIP-E3^63^-induced damage signaling. RPE1 hTERT FlpIn PIP-E3^63^ cells were treated with 1 µg/ml Dox and 5 μM NMS-873 (VCPi) or a corresponding amount of DMSO for 6 h (Dox + VCPi). Alternatively, Dox was added for 12 h and VCP was inhibited for the last 6 h of the treatment (Dox → VCPi). Total ubiquitin levels, checkpoint activation, and E3 expression were analyzed by western blotting of total cell extracts. (**i**) Replication collapse induced by excessive PCNA polyubiquitylation depends on VCP. DNA fiber assays were performed on RPE1 hTERT FlpIn PIP-E3^63^ cells pre-treated for 4 h with 1 µg/ml Dox and 5 μM NMS-873 as indicated. A representative experiment from two replicates is shown. Significance levels were calculated using the two-tailed Mann-Whitney test from the indicated number of fibers per sample (****: p<0.0001). (**j**) Polyubiquitylated PCNA accumulates on chromatin after VCPi. RPE1 hTERT FlpIn PIP-E3^63^ cells were treated with 1 µg/ml Dox and 5 μM NMS-873 or DMSO for 6 h. PCNA was immunoprecipitated from the chromatin fraction, and its ubiquitylation was analyzed by western blotting. (**k**) K63-, K48-, and K11-linked polyubiquitin chains accumulate on PCNA after PIP-E3^63^ expression. Ubiquitin linkages were quantified from the mass spectrometry experiment shown in panel **g** or a corresponding experiment performed after VCPi for 6 h (Extended Data Fig. 5a). The graphs show individual as well as mean values and SD.

Undue replication fork reversal, induced via recruitment of the DNA translocase, ZRANB3, by K63-polyubiquitylated PCNA (*19, 20*), was excluded as a possible cause of the observed replication collapse (Extended Data Fig. 4i). Likewise, two other fork remodelers, HLTF and SMARCAL1 (*30*), as well as known interactors of K63-chains, WRNIP1 and RAP80 (*31, 32*), had no detectable or very mild effects, arguing against their involvement (Extended Data Fig. 4j-l). Depletion of UBE2K, however, suppressed PIP-E3^63^-dependent checkpoint activation (Fig. 2f), thus implicating chain branching in the process. Depletion of HUWE1, an E3 with demonstrated branching activity (*10*), had a milder effect, additive to UBE2K (Fig. 2f). Consistent with a contribution of UBE2K to the action of PIP-E3^63^, co-expression of a catalytically inactive E2 mutant exerted a dominant-negative effect on damage signaling (Extended Data Fig. 4m).

To identify factors mediating the adverse effects induced by polyubiquitylated PCNA, we used SILAC-based mass spectrometry to analyze its interactome by expressing PIP-E3^63^ or its catalytically inactive variant (ΔRING) in HEK 293T cells (Fig. 2g, Extended Data Fig. 5a). In addition to WRNIP1, proteins preferentially associated with polyubiquitylated PCNA included the E3s, HUWE1 and UBR5, a ubiquitin receptor subunit of the proteasome, PSMD4/S5A, as well as the ubiquitin-dependent segregase, VCP. VCP inhibition (VCPi) prevented PIP-E3^63^-induced checkpoint activation and replication fork slowdown (Fig. 2h,i, Extended Data Fig. 5b,c). Under these conditions, polyubiquitylated PCNA, especially its high-molecular weight forms, accumulated on chromatin, with linkages dominated by K63, K48, and to some extent K11 (Fig. 2j,k). Depletion of UBE2K strongly reduced chromatin accumulation of these high-molecular weight forms (Extended Data Fig. 5d), suggesting that VCP dislodges PCNA carrying branched polyubiquitin chains.

Direct evidence for branching on PCNA in cells was obtained by ubiquitin chain restriction (UbiCRest) assays (*33*), employing a K63-specific DUB, AMSH (*34*), on immobilized PCNA isolated from cells expressing PIP-E3^63^ upon VCPi (Fig. 3a). Consistent with K63-polyubiquitylation, AMSH removed nearly all higher ubiquitin conjugates from the immobilized fraction (Fig. 3b). However, the supernatant contained large amounts of oligoubiquitin species of up to four ubiquitin units, indicating branching or extension of the K63-chains to an alternative linkage. Indeed, K48-linked chains were readily detectable by a linkage-specific antibody on the immunoprecipitated PCNA (Fig. 3b). The notion that AMSH almost completely removed them implies that they mostly reside distal to the first K63-linkage. Depletion of UBE2K or co-treatment with a K48-selective DUB, OTUB1 (*35*), reduced the amount of uncleaved oligoubiquitin species in the supernatant, and treatment with a non-selective DUB, USP2cc (*36*), resulted in full disassembly to monoubiquitin (Extended Data Fig. 6a,b). This indicated that the oligomers in the supernatant were mainly linked via K48 and largely assembled by UBE2K, possibly with a contribution of other branching enzymes. Co-expression of a ubiquitin K48R mutant with PIP-E3^63^ strongly stabilized polyubiquitylated PCNA (Extended Data Fig. 6c), while no chains were observed with a K48/63R double mutant. Additional introduction of the K11R mutation did not further stabilize the chains compared to K48R, again confirming the importance of the K48-linkage for PCNA removal.

**Fig. 3.**
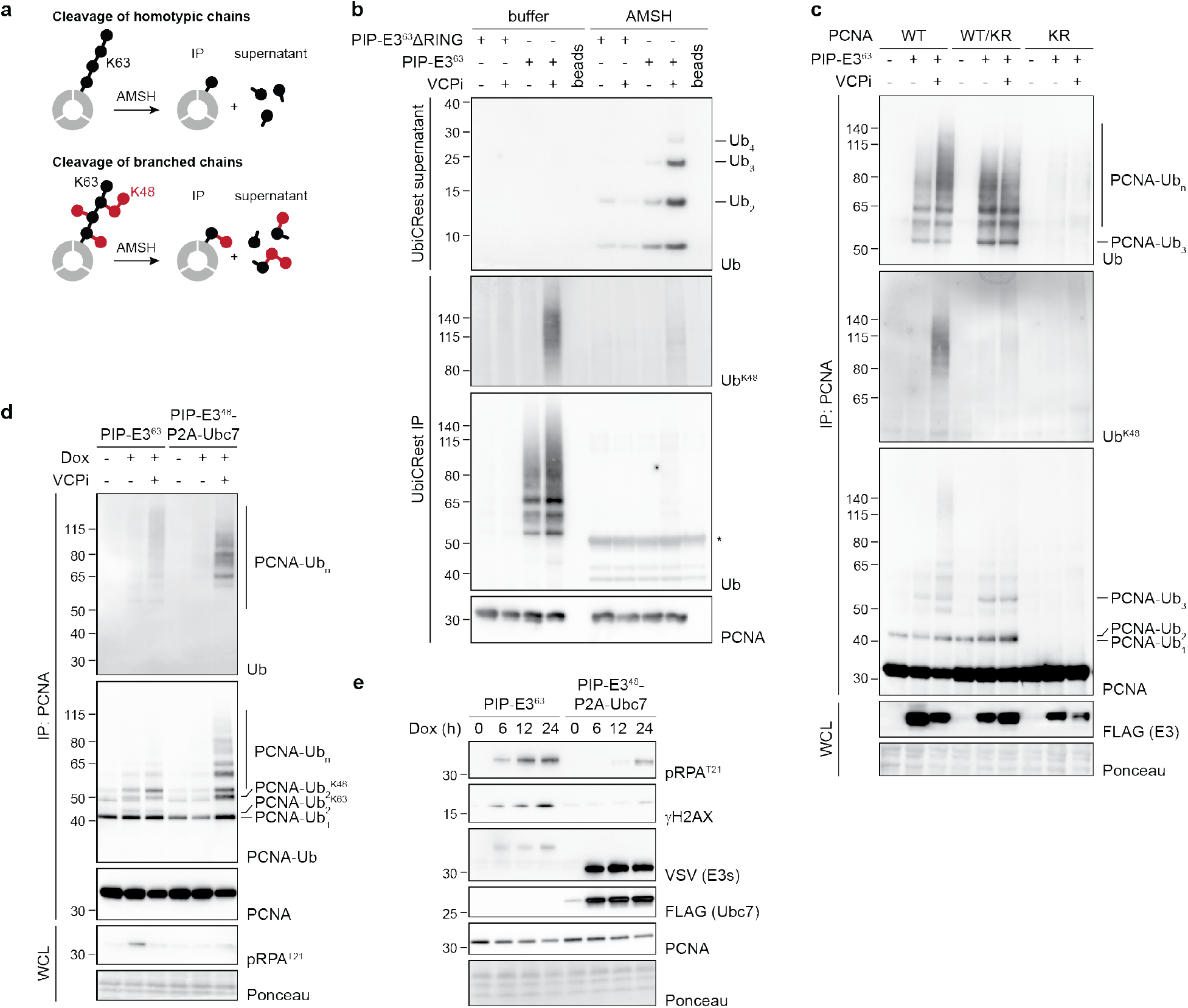
Trans-branching induces PCNA extraction and damage signaling. (**a**) Schematic view of UbiCRest assays on immobilized PCNA carrying either a homotypic K63-linked or a K63-to-K48 branched chain. (**b**) Polyubiquitylated PCNA undergoes K63-to-K48 branching. UbiCRest analysis was performed on PCNA immunoprecipitated from HEK 293T cells transfected with PIP-E3^63^ and treated with 5 μM NMS-873 or DMSO for 6 h. Catalytically inactive E3 (ΔRING) was used as a control. Beads were treated with 3 μM AMSH for 1 h, followed by analysis of supernatant and immobilized fractions (IP) by western blotting. An asterisk indicates cross-reactivity with AMSH. (**c**) Branching requires multiple polyubiquitin chains on the PCNA trimer. PIP-E3^63^ was transiently overexpressed in HEK 293T cells harboring different combinations of wildtype (WT) and mutant (KR) PCNA. Where indicated, VCP was inhibited 3 h prior to cell collection by 5 µM NMS-873. Chromatin-bound PCNA was immunoprecipitated, followed by analysis of ubiquitylation patterns with the indicated antibodies. E3 expression was analyzed in whole cell lysates. (**d**) PIP-E3^48^ affords PCNA polyubiquitylation comparable to PIP-E3^63^ but does not effectively activate DNA damage signaling. RPE1 hTERT FlpIn cells harboring PIP-E3^63^ or PIP-E3^48^-P2A-Ubc7 were allowed to express the enzymes for 6 h with or without VCPi. Ubiquitylation of PCNA was analyzed after PCNA immunoprecipitation from the chromatin fraction and checkpoint activation was detected in whole cell lysates. (**e**) PIP-E3^48^ affords reduced damage signaling compared to PIP-E3^63^. Cells from panel **d** were allowed to express the enzymes for the indicated periods. Checkpoint activation, expression of enzymes, and PCNA levels were analyzed by western blotting in total cell extracts.

Collectively, these data indicate that K63-linked chains on PCNA become further modified by UBE2K with short, predominantly K48-linked chains of two to four ubiquitin units, and that these branched conjugates serve as a signal for VCP-mediated extraction from chromatin and checkpoint activation.

### UBE2K’s trans-branching activity is required for damage signaling upon PCNA extraction

To examine the relevance of UBE2K’s trans-activity, we analyzed PCNA modifications in a panel of HEK 293T-derived cell lines expressing different compositions of the PCNA trimer (*29*): either fully wildtype (WT), exclusively the K164R mutant (KR), or a combination of WT and KR that is expected to result in mixed trimers (WT/KR). Strikingly, despite an overall similar level of PCNA polyubiquitylation in WT and WT/KR cells, stabilization of high-molecular-weight ubiquitin conjugates and accumulation of K48-chains upon PIP-E3^63^ expression and VCPi were observed only in WT, but not WT/KR cells (Fig. 3c). Accordingly, UbiCRest assays revealed reduced branching on PCNA in the WT/KR cell line compared to WT (Extended Data Fig. 6d), and checkpoint signaling was not activated (Extended Data Fig. 6e). These observations suggest that efficient branching and extraction require multiple K63-linked chains on the PCNA trimer, consistent with UBE2K’s *in vitro* properties (Fig. 1e).

If PCNA extraction from chromatin were responsible for the replication collapse induced by PIP-E3^63^, direct modification of PCNA with homotypic K48-linked polyubiquitin chains should circumvent the need for K63-ubiquitylation and likewise result in checkpoint activation. To test this hypothesis, we used a custom PCNA-specific PIP-E3^48^, along with its cognate E2, Ubc7 (*28*), to selectively induce PCNA K48-polyubiquitylation (Extended Data Fig. 6f). As expected, detection of modified PCNA strongly depended on VCPi, indicating that the segregase effectively extracts K48-polyubiquitylated PCNA from chromatin. Side-by-side comparison even revealed a higher total amount of polyubiquitylated PCNA afforded by PIP-E3^48^ compared to PIP-E3^63^ (Fig. 3d). Yet, checkpoint activation by PIP-E3^48^ was much reduced (Fig. 3d,e) and independent of UBE2K (Extended Data Fig. 6g). Thus, PCNA extraction from chromatin per se is unlikely to be the direct cause of the damage response upon unscheduled ubiquitylation of PCNA. Instead, our data suggest that K63-and K48-linked polyubiquitylation need to follow each other in an ordered sequence for efficient activation of checkpoint signaling.

### Branching and VCP-dependent PCNA extraction are induced by DNA replication stress

In untransformed RPE1 hTERT cells, VCPi also resulted in an accumulation of polyubiquitylated PCNA on chromatin after treatment with hydroxyurea (HU), methyl methanesulfonate (MMS) or ultraviolet (UV) radiation (Fig. 4a); and UbiCRest assays revealed branched chains on PCNA in response to HU (Fig. 4b). This demonstrates that branching or extension of K63-linked chains, followed by VCP-dependent extraction from chromatin, is not limited to situations of unscheduled PCNA polyubiquitylation via PIP-E3^63^, but also occurs when the modification is induced by replication stress. Titrations with MMS, HU and camptothecin (CPT) in the absence of VCPi showed the most pronounced effects of UBE2K depletion under high damage loads (Extended Data Fig. 7a-c). To analyze the kinetics of the branching reaction in cells, we performed UbiCRest time courses under VCPi conditions to avoid PCNA extraction and degradation. These revealed a substantial delay in the appearance of K48-branches relative to K63-chains, both when PCNA deubiquitylation was inhibited via USP1i (Fig. 4c) and under conditions where it was allowed (Extended Data Fig. 7d). This delay is consistent with UBE2K’s requirement for the assembly of multiple K63-chains on the PCNA trimer for activity.

**Fig. 4.**
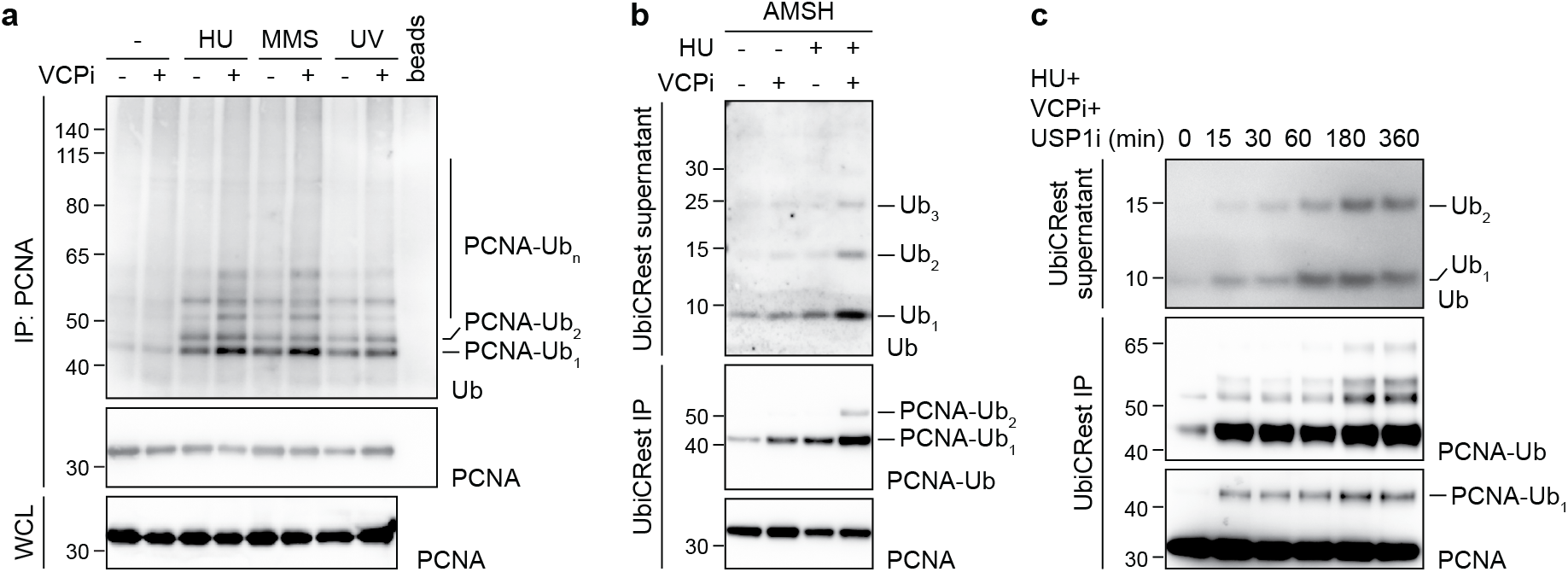
Branching and VCP-dependent PCNA extraction are induced by DNA replication stress. (**a**) VCP removes PCNA from chromatin after DNA damage-induced polyubiquitylation in untransfected cells. RPE1 hTERT cells were treated either with 4 mM HU for 4.5 h or with 0.01% MMS for 1 h, followed by 4.5 h recovery, or with 40 J/m^2^ UV-C irradiation, followed by 4.5 h recovery. Where indicated, VCPi was applied during HU treatment or during recovery from MMS and UV treatment. PCNA was immunoprecipitated from the chromatin fraction and its ubiquitylation was analyzed by western blotting with the indicated antibodies. (**b**) Replication stress induces branching on PCNA. RPE1 hTERT cells were treated with 4 mM HU and VCPi for 2 h. PCNA was immunoprecipitated from the chromatin fraction and subjected to UbiCRest by treatment on beads with 3 µM AMSH for 1 h, followed by analysis of supernatant and immobilized fractions (IP) by western blotting. (**c**) Branching is delayed relative to K63-polyubiquitylation. RPE1 hTERT cells were treated with a combination of 4 mM HU, VCPi and USP1i for the indicated periods. PCNA was immunoprecipitated from the chromatin fraction and subjected to UbiCRest as in Fig. 3b.

We conclude from these results that UBE2K-mediated trans-branching of polyubiquitylated PCNA is a slow, but physiological response to replication fork stalling that becomes important at moderate to high levels of DNA genotoxic stress and induces VCP-mediated extraction of PCNA from chromatin.

### Synthetic lethality between BRCA1 and USP1 deficiency is caused by daughter-strand gaps

We reasoned that excessive K63-polyubiquitylation of PCNA followed by UBE2K-mediated branching and VCP-mediated extraction may underlie not only the damage response and loss of viability upon PIP-E3^63^ expression, but also the lethal effects of USP1i in a BRCA1-deficient background. Consistent with this notion, even short-term VCPi suppressed USP1i-mediated checkpoint activation in BRCA-deficient cells (Fig. 5a) and facilitated the detection of branched polyubiquitin conjugates on PCNA (Fig. 5b). As expected, PCNA degradation, accelerated here by low doses of HU, was also suppressed by VCPi (Fig. 5c, Extended Data Fig. 8a,b). The drop in both chromatin-bound and total PCNA levels suggested that soluble PCNA replenishes the pool removed from chromatin by VCP (Fig. 5c, Extended Data Fig. 8a). As observed with PIP-E3^63^ expression (Fig. 2f), HUWE1 depletion had a minor effect on PCNA degradation and damage signaling when combined with UBE2K depletion (Extended Data Fig. 8c).

**Fig. 5.**
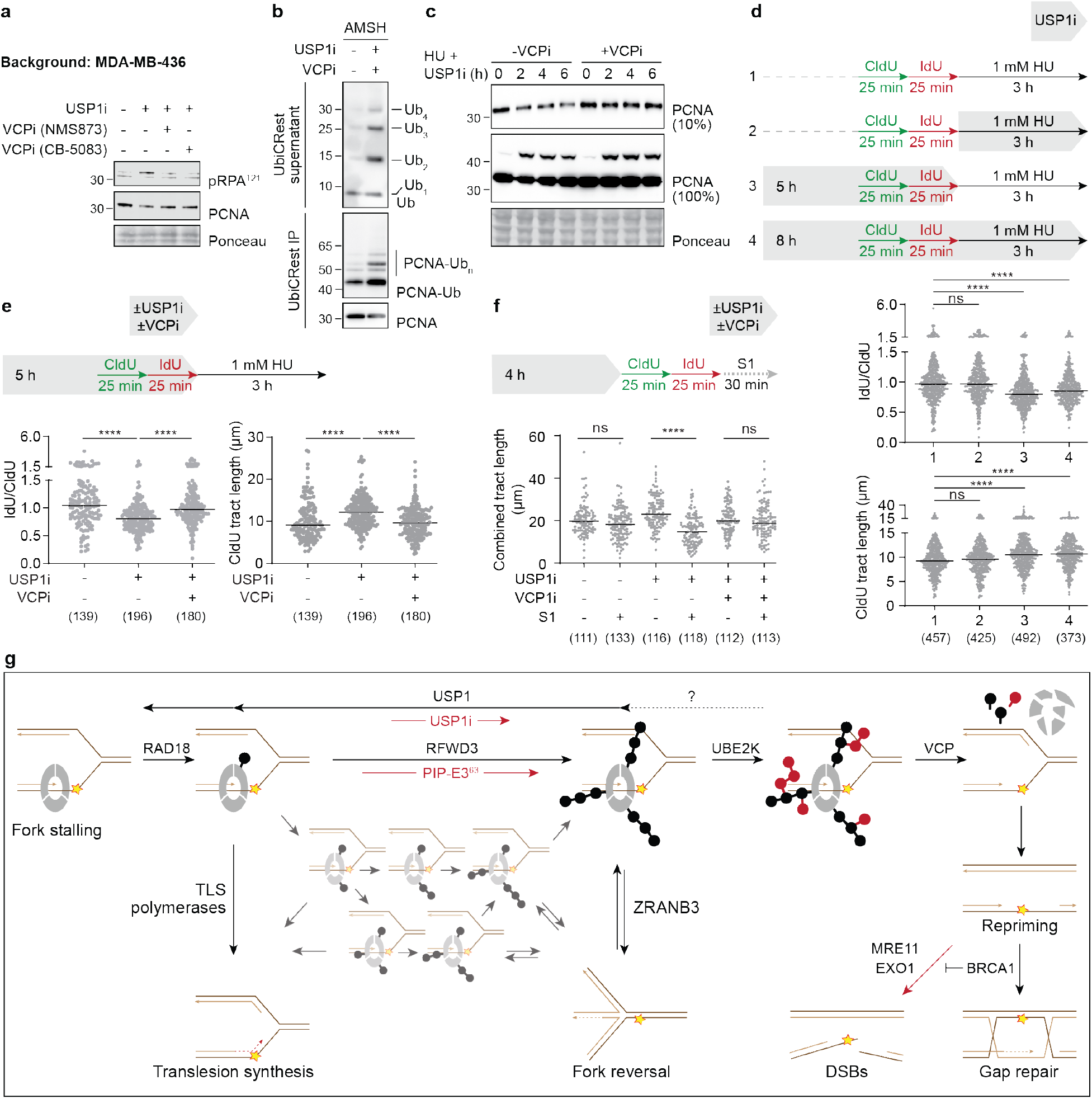
Daughter-strand gaps are causative for toxicity of USP1i in BRCA1-deficient cells. (**a**) Damage signaling in BRCA1-deficient cells depends on VCP. MDA-MB-436 cells were treated with USP1i for 12 h. VCPi was applied as indicated during the final 6 h of the treatment, and checkpoint activation was monitored by western blotting. (**b**) USP1i induces branching on PCNA in BRCA1-deficient cells. UbiCRest assays were performed as in Fig. 3a on MDA-MB-436 cells after USP1i and VCPi for 3 h. (**c**) Total PCNA levels in BRCA1-deficient cells decline upon USP1i. MDA-MB-436 cells were treated with USPi for the indicated periods, and PCNA was analyzed in total cell extracts by western blotting. (**d**) USP1i in BRCA1-deficient cells causes nascent DNA degradation under conditions of replication stress and fork acceleration when applied before, but not during fork stalling. MDA-MB-436 cells were treated as indicated (top), followed by DNA spreading and fiber analysis (bottom). A representative experiment from two replicates is shown. Significance levels were calculated using the two-tailed Mann-Whitney test from the indicated number of fibers per sample (ns: p>0.05; ****: p<0.0001). (**e**) Replication stress-induced nascent DNA degradation and fork acceleration upon USP1i in BRCA1-deficient cells depend on VCP. Fiber assays were performed and analyzed as in panel **d** after the indicated treatments. A representative experiment from two replicates is shown. (**f**) Daughter-strand gaps accumulate in BRCA1-deficient cells upon USP1i in a VCP-dependent manner. Following the indicated treatments, fiber assays were combined with S1 nuclease treatment after cell lysis to reveal gap formation via cleavage of ssDNA. Analysis was as in panel **d**. A representative experiment from two replicates is shown. (**g**) A model illustrating the coordination of replicative DNA damage processing pathways by a complex PCNA-associated ubiquitin code. The scheme shows the gradually growing complexity of ubiquitin conjugates on PCNA as well as major enzymes and readers involved in coordinating the response to replication fork stalling. Red arrows indicate pathological or experimental conditions that drive a DNA damage response due to the hyperaccumulation of daughter-strand gaps.

Taken together, these results suggest that the adverse effects of USP1i in the absence of functional BRCA1 are due to a loss of PCNA following excessive polyubiquitylation involving UBE2K-dependent K63-to-K48 branching. This raised the question as to which stage in the processing of stalled replication forks was sensitive to USP1i. BRCA1 contributes to both the protection of stalled or reversed forks from nucleolytic attack and the repair of postreplicative daughter-strand gaps resulting from fork re-priming downstream of lesions (*37–39*). Lethality upon USP1i could therefore result either from erosion of arrested forks or from failed gap repair. To distinguish between these scenarios, we carried out fork protection assays in combination with USP1i during or prior to HU-induced fork stalling. If the toxic effect of USP1i was due to breakdown of stalled forks, enhanced erosion of nascent DNA should be triggered by USP1i during fork stalling. However, USP1i at this time did not further destabilize replication forks (Fig. 5d, Extended Data Fig. 8d). Instead, addition of the USP1 inhibitor prior to HU treatment enhanced nascent DNA degradation even if the inhibitor was washed out before inducing fork stalling. Likewise, nascent DNA degradation in response to higher doses of HU was intensified specifically by pre-treatment with USP1i, but not by USP1i during fork stalling (Extended Data Fig. 8d). These results are inconsistent with enhanced breakdown of stalled forks in the combined absence of BRCA1 and USP1.

In addition to nascent DNA degradation after pre-treatment with USP1i, we also observed an acceleration of DNA replication (Fig. 5d, Extended Data Fig. 8d) that was suppressed by VCPi (Fig. 5e). Similarly, low-dose HU treatment, often used to analyze the balance between fork reversal and re-priming (*40*), resulted in VCP-dependent lengthening of replication tracts upon USP1i (Extended Data Fig. 8f). As unrestrained replication is a hallmark of daughter-strand gap formation (*38, 40*), we directly examined the presence of daughter-strand gaps by combining the fiber assay with S1 nuclease treatment (*41*). As shown in Fig. 5f, USP1i resulted in enhanced accumulation of daughter-strand gaps, and VCPi reversed this effect. Of note, depletion of PRIMPOL, a factor that contributes to re-priming on the leading strand (*42*), had no effect on damage signaling (Extended Data Fig. 8g). Thus, the gaps responsible for checkpoint activation may predominantly reside on the lagging strand. Alternatively, other mechanisms might drive re-priming of the leading strand under these conditions. In BRCA1-proficient cells, Dox-induced expression of PIP-E3^63^ likewise caused an accumulation of daughter-strand gaps over time (Extended Data Fig. 8h).

To clarify whether the toxicity of PCNA hyper-ubiquitylation was caused directly by the loss of PCNA from chromatin or rather indirectly by the excessive accumulation of daughter-strand gaps, we depleted 53BP1, a damage response mediator that promotes gap formation in BRCA1-deficient cells (*43*). While under these conditions USP1i still efficiently induced PCNA degradation, checkpoint activation was reduced along with enhanced survival (Extended Data Fig. 8i,j), suggesting that gap accumulation rather than loss of PCNA was responsible for the toxicity of USP1i in the BRCA1-deficient background.

We conclude that the massive DNA damage response and subsequent loss of viability, induced either upon USP1i in BRCA1-deficient cells or by PIP-E3^63^ expression, is caused by an excessive accumulation of daughter-strand gaps due to unrestrained re-priming upon VCP-mediated PCNA extraction.

## Discussion

By attributing the toxic effects of excessive PCNA polyubiquitylation to the accumulation of daughter-strand gaps, our findings offer a mechanistic explanation for the synthetic lethality between BRCA1 deficiency and USP1i. This relationship has important implications for the therapy of BRCA1-deficient tumors, particularly as an alternative treatment option in cases of PARP1 inhibitor resistance. At the same time, our study provides insight into ubiquitin signaling by polyubiquitin chain branching. PCNA mono-and K63-polyubiquitylation have served as paradigms for proteasome-independent ubiquitin functions, promoting translesion synthesis and fork reversal, respectively. We now postulate that by inducing VCP-mediated PCNA extraction, K63-to-K48 branching adds a functionality that facilitates re-priming, thereby offering a path to BRCA1-dependent daughter-strand gap repair (Fig. 5g). Thus, branching affords a switching between distinct functionalities of ubiquitin chains. Such scenario calls for a temporal regulation of the branching activity to allow sufficient time for K63-dependent signaling. Importantly, the temporal delay imposed by UBE2K’s trans-activity in combination with USP1-dependent deubiquitylation provides such a timer mechanism, allowing for exploration of alternative, mono-or K63-polyubiquitylation-dependent pathways before committing to re-priming. Our observation that UBE2K’s effects are most pronounced at high damage loads is consistent with this notion of branching-dependent PCNA extraction as a pathway of last resort.

Why then are cells unable to bypass K63-polyubiquitylation and branching by direct K48-polyubiquitylation, which induces PCNA extraction and degradation with equal efficiency? Our data indicate that K63-chains on PCNA have ZRANB3-independent functions beyond their role as seeds for K48-chains. This is consistent with a reported contribution of PCNA polyubiquitylation to chromatin assembly at stalled forks, a CAF-1-dependent pathway that protects forks from nascent DNA degradation (*29*). Additionally, PCNA polyubiquitylation was recently observed in BRCA1-deficient cells to counteract re-priming by re-annealing extensively degraded forks (*44*). We hypothesize that these K63-dependent activities support critical transactions that prepare stalled forks for re-priming and therefore cannot be skipped by direct K48-polyubiquitylation.

The substrate-independent activity of UBE2K suggests that trans-branching may be a general principle to coordinate the hierarchy of ubiquitin signals. In fact, a recent study showed a tendency of UBE2K to extend ubiquitin chains on substrates already carrying multiple ubiquitin units of mixed linkage (*45*). On multimeric substrates or those with multiple ubiquitylation sites, trans-branching therefore has the potential to act as a delayed off-switch for non-proteolytic ubiquitin signals. Although this timer concept has not yet been demonstrated elsewhere, complex functional interactions between K63-and K48-chains suggest such a relationship in other cellular pathways, including chromatin-associated processes (*15*). Future studies will clarify whether other branching E3s also engage in trans-branching or-elongation, and which VCP adaptors are involved in the recognition of specific branched substrates (*8, 46*). Answering these questions will shed light onto a still poorly understood aspect of ubiquitin signaling.

## Methods

### Reagents

Chemicals used in this study were from Merck KGaA, Sigma-Aldrich or Thermo Fisher Scientific unless otherwise stated. Columns and resins used for protein purification were from Cytiva. Antibodies, plasmids, siRNA and bacterial strains and cell lines used in this study are listed in Extended Data Tables 1-4.

### Plasmid construction

Sequences encoding ^VSV^PIP-E3^63^, ^VSV^PIP-E3^48^, and Ubc7 were codon-optimized for use in *H. sapiens*, and a self-cleavable fusion of E3^48^ to Ubc7 was cloned as described (*47*). For transient overexpression and creation of FlpIn cell lines, constructs were cloned into the vector pDEST/FRT/TO-FLAG under control of the CMV promoter, followed by two copies of the TetO sequence. For lentiviral transduction, the sequences encoding ^VSV^PIP-E3^63^, UBE2K (WT or C92A) as well as self-cleavable fusion proteins, ^VSV^PIP-E3^63^-P2A-UBE2K (WT/C92A), were cloned into a pLentiCMV/Tre/3G backbone under control of a minimal CMV promoter and seven repeats of the Tet operator. A sequence encoding His_10_-tagged ubiquitin was placed under the control of the CMV promoter. K-to-R mutations were introduced by site-directed mutagenesis.

A GFP-specific E3^63^ (antiGFP-E3^63-His^) for *in vitro* use was constructed in analogy to the yeast-specific constructs described previously (*28*) by fusing the sequence of the VHH-E domain of an enhancer GFP-specific nanobody (*48*) to the E3^63^ sequence, separated by a 20 amino acid hydrophilic linker. A C-terminal His_6_-tag was appended in a pET24 backbone.

Dimerizable model substrates (^His^FRB-Ub-GFP and ^His^FKBP-Ub-GFP) for *in vitro* use were generated by fusing sequences encoding the respective dimerization domains to a Ub(G76V) sequence, joined by a 20 amino acid hydrophilic linker and a 3C cleavage site, and followed by a C-terminal GFP gene. A His_6_-tag was appended N-terminally in a pET15 backbone.

### Mammalian cell culture

HEK 293T, RPE1 hTERT WT, RPE1 hTERT PCNA(K164R) and MDA-MB-436 cells were cultured in Dulbecco’s Modified Eagle’s Medium (DMEM) supplemented with 2 mM L-glutamine, 10% fetal bovine serum (FBS), 100 U/ml penicillin and 100 μg/ml streptomycin in a humidified atmosphere of 5% CO_2_ at 37°C. Unless specified otherwise, doxycycline, NMS-873 and ML-323 were used at final concentrations of 1 μg/ml, 5 μM and 30 μM, respectively. RPE1 hTERT TREX FlpIn cells were cultured in RPMI medium with the same additives as above. Blasticidin and puromycin were added only after integration of FRT plasmids to final concentrations of 10 μg/ml and 2 μg/ml, respectively.

### Transient transfection

For the transfection of HEK 293T cells, 12 μg of plasmid DNA were mixed with 40 μl of 1 mg/ml polyethyleneimine (Polysciences) in 2 ml of serum-free DMEM, vortexed and incubated for 10 min at room temperature. The transfection mix was added to 5·10^6^ cells. After 4 h in the cell culture incubator, the medium was exchanged to full DMEM. Cells were collected 24 h after transfection. For overexpression of His-tagged ubiquitin, 2.5·10^6^ cells were used, and cells were collected 48 h after transfection. RPE1 hTERT cells were transfected with Fugene HD reagent according to the manufacturer’s instructions using 3 μg of plasmid DNA and 9 μl of the transfection reagent per 2·10^6^ cells.

Transfections of siRNAs (Extended Data Table 3) were performed with Lipofectamine RNAiMAX reagent according to the manufacturer’s instructions. Briefly, 10 μl of 10 μM siRNA solution were mixed with 25 μl of transfection reagent in 1 ml Opti-MEM^®^ Reduced Serum Medium (Life Technologies), incubated for 15 min at room temperature and added to cells together with 4 ml Opti-MEM. The transfection mix was exchanged for full medium after 4 h and cells were analyzed 72 h after transfection.

### Generation of stable cell lines

For generation of FlpIn cell lines, vectors harboring the target constructs were mixed with the Flp-recombinase expression plasmid, pOG44, in a mass ratio of 1:9 and used for transfection of RPE1 hTERT TREX FlpIn cells with Fugene HD reagent according to the manufacturer’s instructions. 48 h after transfection, blasticidin and puromycin (Invivogen) were added to the medium to final concentrations of 10 μg/ml and 2 μg/ml, respectively. The medium was exchanged every 2-3 days until colonies of the desired cell line appeared.

Generation of non-FlpIn stable cell lines was carried out by lentiviral transduction. Lentivirus particles were produced by transfecting HEK 293T cells with packaging vectors (pMDLg/pRRE, pRSV-Rev, pMD2.G) and a vector encoding PIP-E3^63^ or tetracycline transactivator rtTA3 in a mass ratio of 1:1:1:1.6 with polyethyleneimine as described above. 24 h after transfection, supernatants containing lentiviral particles carrying PIP-E3^63^ or rtTA3 DNA were combined and passed through a 0.45 μm syringe filter. Polybrene was added to a final concentration of 5 μg/mL, and viral supernatants were added to 5·10^5^ (for RPE1 hTERT) or 2·10^6^ (for HEK 293T) recipient cells. Medium was exchanged the next day, and blasticidin and G418 were added 48 h after transduction to final concentrations of 10 μg/ml and 400 μg/ml, respectively. Medium was exchanged every 2-3 days for at least two weeks until only antibiotic-resistant cells remained in the culture.

### Flow cytometry

10^6^ RPE1 hTERT FlpIn cells harboring relevant integrated constructs were plated on 10 cm dishes and treated with 1 µg/ml doxycycline for 24 or 48 h. Cells were rinsed once with PBS and trypsinized in 1 ml of 0.05% trypsin-EDTA at 37°C for 5 min. Detached cells were diluted in 9 ml of full DMEM, centrifuged for 3 min at 300·g, washed once with 1 ml of ice-cold PBS and centrifuged again. The supernatant was discarded, and cells were fixed by drop-wise addition of 1 ml of 70% ethanol pre-cooled to-20°C upon constant vortexing. Cell suspensions were incubated at 4°C overnight and, if needed, further stored at-20°C. For propidium iodide (PI) staining, cells were centrifuged for 10 min at 1,000·g, washed once with 1 ml of ice-cold PBS, and centrifuged again for 10 min at 1,000·g. PBS was removed, and cell pellets were resuspended in 500 µl of staining buffer (100 µg/ml PI and 20 µg/ml RNase A in PBS), followed by incubation for 30 min at room temperature in the dark. For cell cycle analysis of siRNA-transfected cells, transfections were performed with Lipofectamine RNAiMAX as described above, doxycycline was added to cells for 48 h, and cells were collected 72 h after transfection.

Data were acquired on a LSRFortessa Cell Analyzer (BD Biosciences). Data analysis was performed with FlowJo (BD Biosciences). The population of RPE1 hTERT cells was gated using FSC-A x SSC-A parameters, and a single-cell population was selected (YG561nm 610/20 BP W x A). DNA content was visualized in histogram plots (YG561nm 610/20 BP channel) that were normalized to the maximum signal for direct comparison of samples.

For 2D analyses, cells were treated with 10 µM EdU 30 min prior to collection and fixation as described above. Click reactions were performed by incubating the cell suspension in Click-iT buffer (5 µM Alexa Fluor^®^ 647 picolyl azide, 2 mM copper (II) sulfate and 100 mM sodium ascorbate in PBS) for 30 min in the dark, followed by three washes in PBS and PI staining as described above. The EdU-Alexa Fluor 647 signal was detected in the RL 640nm 670/30 BP channel. A minimum of 10,000 cells per sample were analyzed.

### Viability and colony formation assay

For cell viability assays, 2·10^3^ MDA-MB-436 cells were seeded in 96-well plate in triplicates, allowed to adhere overnight, followed by the addition of ML-323 or doxycycline. After 72 h, 20 μl of CellTiter-Blue Reagent (Promega) were added per 100 μl of medium and incubated for 4 h at 37°C. Fluorescence was recorded with a Tecan Spark 20M microplate reader (excitation filter: 535/20 nm; emission filter: 595/35 nm), and intensity values were normalized to the untreated condition. If required, siRNA transfection was performed 24 h prior to the plating in 96-well dishes.

For colony formation assays, 300 cells were seeded on 10 cm dishes in triplicate and allowed to adhere overnight, followed by addition of doxycycline to a final concentration of 1 µg/ml. After 10 days, the medium was removed, colonies were fixed on the plate with a 3:1 (v/v) mixture of methanol and glacial acetic acid for 10 min at room temperature, followed by incubation with 1% aqueous crystal violet solution for 10 min at room temperature. Subsequently, dishes were washed with distilled water until the background was clear, air-dried overnight at room temperature and imaged with an Epson Perfection V700 scanner. The total area covered by the cells was quantified using the ImageJ software (NIH) and results were normalized to untreated cells for each cell line.

### Preparation of whole cell lysates

Cells were collected in ice-cold PBS and centrifuged for 3 min at 1,000·g. The supernatant was removed, and cell pellets briefly frozen in liquid nitrogen. Subsequently, cells were lysed in RIPA buffer (50 mM Tris-HCl, pH 7.5, 1% IGEPAL, 0.5% sodium deoxycholate, 0.1% SDS, 150 mM NaCl, 2.5 mM MgCl_2_) supplemented with cOmplete™ protease inhibitor cocktail (Roche) and 0.625 U/μl Sm Nuclease (produced in-house) for 30 min on ice. Samples were centrifuged at 21,100·g for 10 min, and the supernatant was transferred to a new pre-cooled tube. Protein concentration was normalized using the Bio-Rad Protein Assay (Bio-Rad), and samples were mixed 3:1 with 4x NuPAGE LDS sample buffer and incubated at 95°C for 5 min.

### Western blotting

Proteins were separated by electrophoresis on gradient 4-12% Bis-Tris polyacrylamide gels in 1x MOPS buffer (50 mM MOPS, 50 mM Tris, 0,1 % SDS, 1 mM EDTA). For the detection of monoubiquitin and low-molecular weight ubiquitin chains, gels were run in 1x MES buffer (50 mM MES hydrate, 50 mM Tris, 0.1% SDS, 1mM EDTA). Proteins were transferred onto nitrocellulose membranes using a Trans-blot Turbo transfer system (Bio-Rad). Membranes were blocked in 5% milk in PBST and incubated with primary antibodies overnight at 4°C. Washes were performed with PBST and incubation with secondary antibodies was for 1 h at room temperature. After renewed washing, membranes treated with HRP-conjugated secondary antibody were developed using the Amersham ECL Prime or ECL Select Western Blotting Detection Reagent (Cytiva) according to the manufacturer’s instructions. Chemiluminescence signals were recorded with a Fusion FX (Vilber) instrument. Membranes treated with IRDye-coupled secondary antibodies were imaged using an Odyssey® CLx (LI-COR) instrument.

When VU1 monoclonal anti-ubiquitin antibody was used, membranes after transfer were rinsed twice in PBS and incubated in 0.5% aqueous glutaraldehyde solution for 20 min at room temperature. Membranes were then blocked for 2 h in 5% milk solution in PBST, and incubation with antibodies and development of the HRP signal were performed as described above.

### Detection of PCNA ubiquitylation in chromatin

5·10^6^ cells per sample expressing PIP-E3s or treated with DNA damaging agents were collected in ice-cold PBS and centrifuged at 1,000·g for 3 min. The supernatant was discarded, and the pellet resuspended in 1 ml of 10 mM HEPES pH 7.5,10 mM KCl, 1,5 mM MgCl_2_, 0,35 M sucrose, 10% glycerol, 10 mM N-ethylmaleimide, and 0.1% Triton-X100. After incubation for 10 min on ice and centrifugation at 1,300·g for 5 min, the pellet was frozen in liquid nitrogen to facilitate the disintegration of nuclei. The pellet was then resuspended in 1 ml RIPA buffer supplemented with 10 mM N-ethylmaleimide, cOmplete™ protease inhibitor cocktail, and 0.625 U/μl Sm Nuclease, and incubated for 30 min on ice, followed by addition of 60 μl of 5 M NaCl and a further 5 min of incubation on ice. Lysates were cleared by centrifugation at 21,100·g for 10 min, followed by addition of 1.5 μg of antibody PC10 and incubation at 4°C for 1 h with agitation. Protein A agarose beads (40 μl of a 50% slurry) were washed once with 1 ml RIPA buffer and added to cell lysates. After overnight incubation, beads were washed three times with 1 ml RIPA buffer each and the supernatant was removed completely after the last wash. Proteins were eluted from beads in 50 μl of 2x NuPAGE LDS buffer supplemented with 20 mM DTT for 15 min at 95°C and subjected to SDS polyacrylamide gel electrophoresis and western blotting. To avoid cross-reactivity with the mouse monoclonal PC10 antibody, rabbit antibodies against PCNA, PCNA-Ub or ubiquitin were used for blotting.

### UbiCRest assays

HEK 293T and RPE1 hTERT cells were transfected as described above and treated for 6 h with 5 μM NMS-873. For detection of chain branching in the MDA-MB-436 cell line, cells were simultaneously treated with 5 μM NMS-873 and 30 μM ML-323 for 3 h. After cell collection and chromatin fractionation, PCNA was immunoprecipitated from chromatin as described in the section above. N-ethylmaleimide was avoided throughout the protocol. After 3 washes with RIPA buffer, beads were washed twice with 1x Ubi buffer (20 mM HEPES, 100 mM NaCl, 0.1% Tween 20, 5 mM MgCl_2_) and resuspended in 80 μl of 1x Ubi buffer supplemented with 10 mM DTT. Deubiquitylating enzymes AMSH, OTUB1, and/or ^His^USP2cc were then added to final concentrations of 3 μM, 10 μM and 1 μM, respectively. Beads were incubated at 37°C for 1 h with constant shaking and additionally manually vortexed every 10 min. Beads were centrifuged for 3 min at 1,000·g, and 60 μl of the supernatant were mixed with 20 μl of 4x LDS supplemented with 40 mM DTT, followed by incubation at 95°C for 5 min and analysis by western blotting with VU1 anti-ubiquitin antibody (‘UbiCRest supernatant’). The remaining supernatant was completely removed from the beads with the help of a syringe, and beads were resuspended in 50 μl of 2x LDS supplemented with 20 mM DTT. Beads were incubated at 95°C for 15 min, centrifuged for 3 min at 1,000·g, and eluted proteins were analyzed by western blotting against ubiquitin, Ub^K48^ or PCNA (‘UbiCRest IP’).

### Denaturing Ni-NTA pulldown

2.5·10^6^ HEK 293T cells per condition were plated on 10 cm dishes, transfected with PIP-E3^63^ and respective His-tagged ubiquitin mutants and allowed to express the proteins for 48 h. Cells were briefly washed on plates and collected in cold PBS. 10% of the cell pellet were used to prepare whole cell lysates. For the pulldown, the remaining 90% of the pellet were resuspended in 1.5 ml buffer A (6 M Guanidine-HCl, 50 mM NaH_2_PO_4_, 50 mM Na_2_HPO_4_, 10 mM Tris-HCl pH 8.0, 0.1% Tween 20) and briefly sonicated. Imidazole was added to a final concentration of 15 mM, and the lysate was incubated with 40 μl of 50% Ni-NTA slurry (Qiagen) overnight at 4°C with agitation. The beads were then washed twice with 1 ml buffer A and 4 times with 1 ml buffer B (8 M urea, 80 mM NaH_2_PO_4_, 20 mM Na_2_HPO_4_, 10 mM Tris-HCl pH 6.3, 0.1% Tween 20). After the last wash, the supernatant was completely removed with the help of a syringe, and proteins were eluted in 50 μl elution buffer (2x LDS, 200 mM imidazole) at 95°C for 10 min and analyzed by western blotting as described above.

### DNA fiber assays

Cells were treated with 50 µM CldU for 25 min, briefly washed 3 times with pre-warmed PBS, and treated with 50 µM IdU. For replication stress conditions, 0.5 mM HU were added during IdU labeling, and incubation was for 120 min. For fork protection assays, 1 mM or 4 mM HU were added to cells after a 25 min IdU pulse for 3 or 5 h, respectively. Subsequently, cells were washed 3 times with ice-cold PBS, trypsinized for 10 min at 37°C, supplemented with DMEM and centrifuged at 300·g for 5 min. Cell pellets were resuspended in PBS at 1.75·10^5^ cells/ml and the suspensions were kept on ice. At this stage, samples were blinded.

For S1 nuclease treatment, cells were washed once with PBS after the CldU/IdU labeling and permeabilized with CSK buffer (25 mM HEPES, 1 mM EDTA, 50 mM NaCl, 3 mM MgCl_2_, 300 mM sucrose, 0.5% Triton-X100) directly on the plate for 8 min at room temperature, followed by treatment with 20 U/ml S1 nuclease in S1 buffer (30 mM sodium acetate, 10 mM ZnSO_4_, 5% glycerol, 50 mM NaCl) for 30 min at 37°C. Cells were then collected in 0.1% BSA in PBS, centrifuged at 1,000·g for 3 min, and resuspended in 0.1% BSA in PBS at a concentration of 1.75·10^5^ cells/ml.

For DNA spreading, 7.5 µl of lysis buffer (200 mM Tris-HCl pH 7.5, 50 mM EDTA, 0.5% SDS) were mixed with 4 µl of cell suspension on the surface of SuperFrost Plus™ Gold Adhesion Microscope Slides (Epredia) and incubated for 9 min at room temperature. Slides were then tilted to allow the drop to run down the slide, and air-dried. Thereafter, slides were fixed overnight in a 3:1 (v/v) mix of methanol and acetic acid at 4°C. After washing twice for 3 min in PBS, the DNA was denatured in 2.5 M HCl for 90 min at room temperature, followed by six washes in PBS for 2.5 min each. The spreads were then blocked for 40 min at room temperature in blocking buffer (2% BSA in PBST, sterile filtered), followed by a 2.5 h incubation with primary antibodies (mouse anti-BrdU, 1:100, and rat anti-BrdU, 1:100). After three washes of 5 min each with PBST, slides were incubated with secondary antibodies (goat anti-mouse Alexa Fluor 647, 1:500, and goat anti-rat Alexa Fluor 488, 1:500). Slides were then washed three times for 5 min in PBST, once in distilled water, completely air-dried in the dark and mounted with Prolong Gold AntiFade Mountant. Images of DNA fibers were acquired with a Widefield Fluorescence Microscope (Thunder, LASX software, Leica) (HC PL APO CS2 63x/1.40 oil immersion objective; LED illumination and the corresponding emission filters: 635 nm, 642/80 and 475 nm, 535/70). At least 10 different fields per slide were imaged, and a minimum of 100 fibers were quantified per condition. Tract lengths were quantified using Fiji/ImageJ.

### Mass spectrometry sample preparation

For SILAC labeling, HEK 293T cells were cultured in SILAC DMEM supplemented with dialyzed FBS and either L-lysine and L-arginine (light), L-lysine [^2^H_4_] and L-arginine [^13^C_6_] (medium, Cambridge Isotope Laboratories) or L-lysine [^13^C_6_, ^15^N_2_] and L-arginine [^13^C_6_, ^15^N_4_] (heavy, Cambridge Isotope Laboratories) for a minimum of two weeks. For analysis of the PCNA interactome without VCPi, 20·10^6^ cells per condition were plated in triplicates, transfected the next day with PIP-E3^63^ or PIP-E3^63^ΔRING for 24 h, followed by immuno-precipitation of PCNA from chromatin as described above. For two of the replicates, PIP-E3^63^ was expressed in ‘heavy’-and PIP-E3^63^ΔRING in ‘light’-labeled cells, whereas for the third replicate a label switch was performed. The experiment involving VCP inhibition was performed analogously, with the exception that three conditions were compared (PIP-E3^63^ΔRING + VCPi; PIP-E3^63^ + DMSO; PIP-E3^63^+ VCPi) and cells were treated with 5 μM NMS-873 or the corresponding amount of DMSO 6 h prior to cell collection. The beads from all SILAC conditions were combined during the last wash. Bound proteins were eluted in 2x NuPAGE LDS Sample Buffer supplemented with 1 mM DTT, heated at 95°C for 15 min, alkylated with 5.5 mM chloroacetamide for 30 min at room temperature, and separated on 4-12% gradient Bis-Tris gels. Proteins were stained using a Colloidal Blue Staining Kit (Life Technologies) and digested in-gel(*49*) using MS-grade trypsin (Serva). Peptides were extracted from the gel using a series of increasing acetonitrile percentage and desalted using reversed-phase C18 StageTips (*49*).

### Mass spectrometry data analysis

Samples were analyzed on a quadrupole Orbitrap mass spectrometer (Exploris 480, Thermo Scientific) equipped with a UHPLC system (EASY-nLC 1,200, Thermo Scientific). They were loaded onto a C18 reversed-phase column (55 cm length, 75 mm inner diameter) and eluted with a gradient from 2.4 to 32% acetonitrile containing 0.1% formic acid in 90 min. The mass spectrometer was operated in data-dependent mode, automatically switching between MS and MS2 acquisition. Survey full-scan MS spectra (m/z 325–1,300, resolution: 60,000, target value: 300%, maximum injection time: 28 ms) were acquired in the Orbitrap. The 15 most intense precursor ions were sequentially isolated, fragmented by higher energy C-trap dissociation (HCD), and scanned in the Orbitrap mass analyzer (normalized collision energy: 30%, resolution: 15,000, target value: 100%, maximum injection time: 40 ms, isolation window:

1.4 m/z). Precursor ions with unassigned charge states, as well as with charge states of +1 or higher than +6, were excluded from fragmentation. Precursor ions already selected for fragmentation were dynamically excluded for 25 s. Raw data files were analyzed using MaxQuant (*50*). Parent ion and MS2 spectra were searched against a reference proteome database containing human protein sequences obtained from UniProtKB (version 2020_02) using the Andromeda search engine (*51*). Spectra were searched with a strict trypsin specificity and allowing up to two miscleavages. Cysteine carbamidomethylation was searched as a fixed modification, whereas ubiquitin remnants (di-glycine-lysine), protein N-terminal acetylation, methionine oxidation, and N-ethylmaleimide modification of cysteines (mass difference to cysteine carbamidomethylation) were searched as variable modifications. The dataset was filtered based on posterior error probability (PEP) to arrive at a false discovery rate of below 1% estimated using a target-decoy approach (*52*). Statistical analysis and MS data visualization were performed using the R software environment. P-values and false discovery rates were calculated using a moderated t test (Limma algorithm) (*53*). A list of all identified proteins with enrichments and p-values for each experiment is provided as an Extended Data Excel Table.

### Recombinant protein production

All recombinant proteins were produced in *Escherichia coli*. Ubiquitin variants, ^His^Uba1, ^His^Ubc13, ^GST^Ubc13, Mms2, UBCH5c(S22R), ^GST^PIP-E3^63^, ^His^PCNA, ^His^USP2cc, OTUB1, and

AMSH were purified as described (*28, 33, 54–59*) with minor modifications. ^His^UBE2K was purified by immobilized metal affinity chromatography (IMAC) followed by size exclusion chromatography (SEC) in storage buffer (50 mM HEPES pH 7.4, 150 mM NaCl, 1 mM DTT, 10% glycerol). His_6_-tagged antiGFP-E3^63^ was produced in *E. coli* BL21(DE3)-RIL LOBSTR (Kerafast) grown in TB medium supplemented with 100 µM ZnCl_2_ and purified by IMAC, followed by SEC in 20 mM HEPES pH 7.4, 150 mM NaCl, 5% glycerol. ^His^FRB-Ub-GFP and ^His^FKBP-Ub-GFP were produced in *E. coli* BL21(DE3)-RIL LOBSTR grown in TB medium, purified by IMAC, and transferred into storage buffer (20 mM HEPES pH 8.0, 150 mM NaCl, 5% glycerol).

Mixed trimers of human PCNA were produced by co-expression of ^MBP-3C^PCNA(K164R) and ^His^PCNA from a single, bicistronic plasmid in *E. coli*. Cells were lysed in IMAC A buffer (50 mM Tris-HCl pH 8.0, 300 mM NaCl, 15 mM imidazole) supplemented with SIGMAFAST protease inhibitor cocktail, 2 mM MgCl_2_, and 50 U/ml SmNuclease by two passes through a high-pressure cell disruptor (Constant Systems) in continuous flow at 4°C. Triton X-100 was added to 0.1%, followed by incubation for 20 min at 4°C and clearance of the lysate by centrifugation at 40,000·g for 20 min at 4°C. The cleared lysate was loaded onto a HisTrap HP 5 ml column using an NGC chromatography system (Bio-Rad). Bound protein was eluted in a multi-step gradient using IMAC B buffer (50 mM Tris-HCl pH 8.0, 300 mM NaCl, 250 mM imidazole), thereby separating complexes containing different ratios of ^His^PCNA and ^MBP-3C^PCNA(K164R). Relevant complexes were pooled and applied separately to a Superdex 200 16/600 pg size exclusion column equilibrated in 50 mM HEPES pH 7.4, 150 mM NaCl. Trimers harboring one or two mutant subunits eluted at different volumes according to the size difference introduced by the MBP-tag on ^MBP-3C^PCNA(K164R). The MBP-tag was subsequently cleaved by GST-3C protease overnight at 4°C. Mixed trimers containing ^His^PCNA and PCNA(K164R) were then purified from the cleavage reaction by IMAC using a HisTrap HP 5 ml column, followed by SEC on a Superdex 200 Increase 10/300 column (50 mM HEPES pH 7.4, 200 mM NaCl, 1 mM DTT, 10% glycerol). The final complexes eluted at a volume reflecting trimeric PCNA.

### In vitro ubiquitylation of PCNA

PCNA was monoubiquitylated by UBCH5c(S22R) and purified as described (*55, 58*). This was used as a substrate to prepare K63-linked poly-or diubiquitylated PCNA in reactions containing 1 µM monoubiquitylated PCNA, 5 µM ubiquitin, 50 nM ^His^Uba1, 0.5 µM ^His^Ubc13·Mms2 (E2), 1 µM ^GST^PIP-E3^63^ and 100 µM ATP in ubiquitylation buffer (40 mM HEPES pH 7.4, 50 mM NaCl, 8 mM magnesium acetate) by incubation for 30 or 40 min at 30°C. The products of these reactions served as input for UBE2K activity assays.

To compare the activity of UBE2K towards unmodified, monoubiquitylated, and K63-polyubiquitylated PCNA, reactions were set up in ubiquitylation buffer either with purified unmodified or monoubiquitylated PCNA or with the unpurified products of a K63-polyubiquitylation reaction, using 1.5 µM UBE2K. Reactions were incubated at 37°C for 3 h and products were analyzed by western blotting against Ub^K48^, PCNA, and UBE2K.

To examine branching on fully diubiquitylated PCNA carrying the K48R mutation at the proximal and/or distal positions, PCNA was first monoubiquitylated using either WT Ub or Ub(K48R). After purification, a single K63-linkage was generated on each subunit using either

Ub(K63R) or Ub(K48/63R) in a polyubiquitylation reaction with ^His^Ubc13·Mms2 and ^GST^PIP-E3^63^ as described above. Activity of UBE2K towards these preparations of fully diubiquitylated PCNA was then tested as described above using 5 µM Ub(K48/63R).

To examine branching on diubiquitylated PCNA carrying either a single or up to three diubiquitin units, a preparation of either mixed trimers (prepared as described above) or WT ^His^PCNA was monoubiquitylated using Ub^WT^. After purification, the monoubiquitylated PCNA trimers (1 µM) were further ubiquitylated by a single K63-linkage as described above but using 20 µM of Ub(K63R). Without further purification, an equal volume of reaction buffer containing 100 nM of ^His^Uba1, 3 µM UBE2K, 20 µM Ub(K63R), and 200 µM ATP was added and the reactions were incubated at 37°C. Reactions were terminated at the indicated time points by addition of 4x LDS supplemented with 40 mM DTT, denatured for 3 min at 95°C, and analyzed by western blotting against PCNA and Ub^K48^.

### In vitro ubiquitylation of Ub-GFP

*In vitro* branching on model substrates was analyzed in a two-step procedure. First, preparative reactions to produce K63-diubiquitylated FRB-Ub-GFP and FKBP-Ub-GFP were performed. Each protein was modified separately in reactions containing 4 µM substrate, 50 nM E1, 0.2 µM Ubc13·Mms2 (E2), 2 µM antiGFP-E3^63^, 100 µM ATP, and 4 µM Ub(K63R) in ubiquitylation buffer for 60 min at 30°C. Then, the K63-diubiquitylated substrates were combined, supplemented with an additional 100 µM ATP and 4 µM Ub(K63R), and incubated with 2 µM UBE2K and 5 µM rapamycin (Biozol) where indicated. Branching reactions were performed for 30 and 60 min at 37°C. Reactions were terminated by addition of 4x LDS supplemented with 40 mM DTT, incubated for 2 min at 95°C, and analyzed by western blotting against Ub^K48^, FRB, FKBP, and UBE2K. For the asymmetric combination, the Ubc13·Mms2-dependent ubiquitylation step was omitted for one of the substrates, and K63-ubiquitylation of the other substrate was carried out with ^GST^Ubc13. Prior to combining the two substrates, ^GST^Ubc13 was removed by passage over glutathione beads to prevent further K63-ubiquitylation.

### AlphaFold 3-based in silico model building

Mimics of polyubiquitylated PCNA were generated *in silico* following an approach used previously to solve the structure of monoubiquitylated yeast PCNA (*60*). A di-ubiquitinated human split PCNA (^spl^PCNA) sequence was generated by splitting the PCNA sequence into an unmodified N-terminal part (^spl^PCNA^N^, aa 1-163) and a C-terminal portion (^spl^PCNA^C^, aa 165-261) extended at its N-terminus by two copies of the ubiquitin sequence fused head-to-tail (Ub-Ub) as an approximation of the K63-linkage. To mimic the isopeptide bond between ubiquitin and PCNA, two glycine residues were inserted between Ub_2_ and ^spl^PCNA^C^ in place of K164. The AlphaFold 3 Server (*27*) was queried with three copies of ^spl^PCNA^N^ and Ub-Ub-^spl^PCNA^C^ each, one copy of UBE2K and one human monoubiquitin (Ub) to provide a potential donor ubiquitin. A 220 bp fragment of the coding sequence of human mTOR (NCBI: NM_004958, positions 6182 to 6401) was chosen arbitrarily and added as a generic DNA scaffold. A complementary strand was generated in the query by AlphaFold 3. Using automatic seed selection, AlphaFold 3 randomly selected seed 1527108903. The resulting structure predictions were visualized using PyMol and the predicted alignment error (PAE) matrix for each model was generated using the PAE Viewer web tool from the University of Göttingen (*61*). Model #2 was selected as a potential representation of the trans-branching event. The sequences used for model generation are listed in Extended Data Table 5.

## Data availability

The MS-based proteomics data have been deposited with the ProteomeXchange Consortium via the PRIDE partner repository with the dataset identifier PXD054064. All other data are available in the main text or the supplementary materials.

Reviewer access details:

Log in to the PRIDE website using the following details:

Project accession: PXD054064

Token: yErUg3PCEMF2

Alternatively, reviewers can access the dataset by logging in to the PRIDE website using the following account details:

Username: Password: YhMv8KjzYDhh

## Acknowledgments

We thank Alexandra Belayew, Anja Bielinsky, Simon Boulton, Lucian Moldovan, and Svend Petersen-Mahrt for reagents, Jiaxuan Chen and Ronald Wong for help with experiments, IMB’s Core Facilities for Microscopy, Flow Cytometry, Protein Production, Proteomics, and the Media Lab for their services and reagents, and Cindy Meister, Hans-Peter Wollscheid, and Nicola Zilio for critical reading of the manuscript. This research was funded by the Deutsche Forschungsgemeinschaft (DFG, German Research Foundation) – Project-ID 393547839 (SFB 1361) to PB and HDU, and Project-ID BE 5342/2-1 (FOR 2800) to PB, and by an ERC Advanced Grant (#) to HDU. Funding from the German Research Foundation also supported the LSRFortessa (#210253511, IMB Flow Cytometry Core Facility) and the Exploris 480 (#393547839, IMB Proteomics Core Facility).

## Author contributions

KP and HDU conceptualized the study and wrote the initial draft; KP, CR, MSS, IM, NCK, and AT performed experiments and analyzed data; PB and HDU acquired funding and provided supervision; KP, CR, IM, PB, and HDU contributed to reviewing and editing of the manuscript.

## Materials & Correspondence

All materials used in the analysis will be made available by the corresponding author upon request.

---

**Extended Data Fig. 1.**
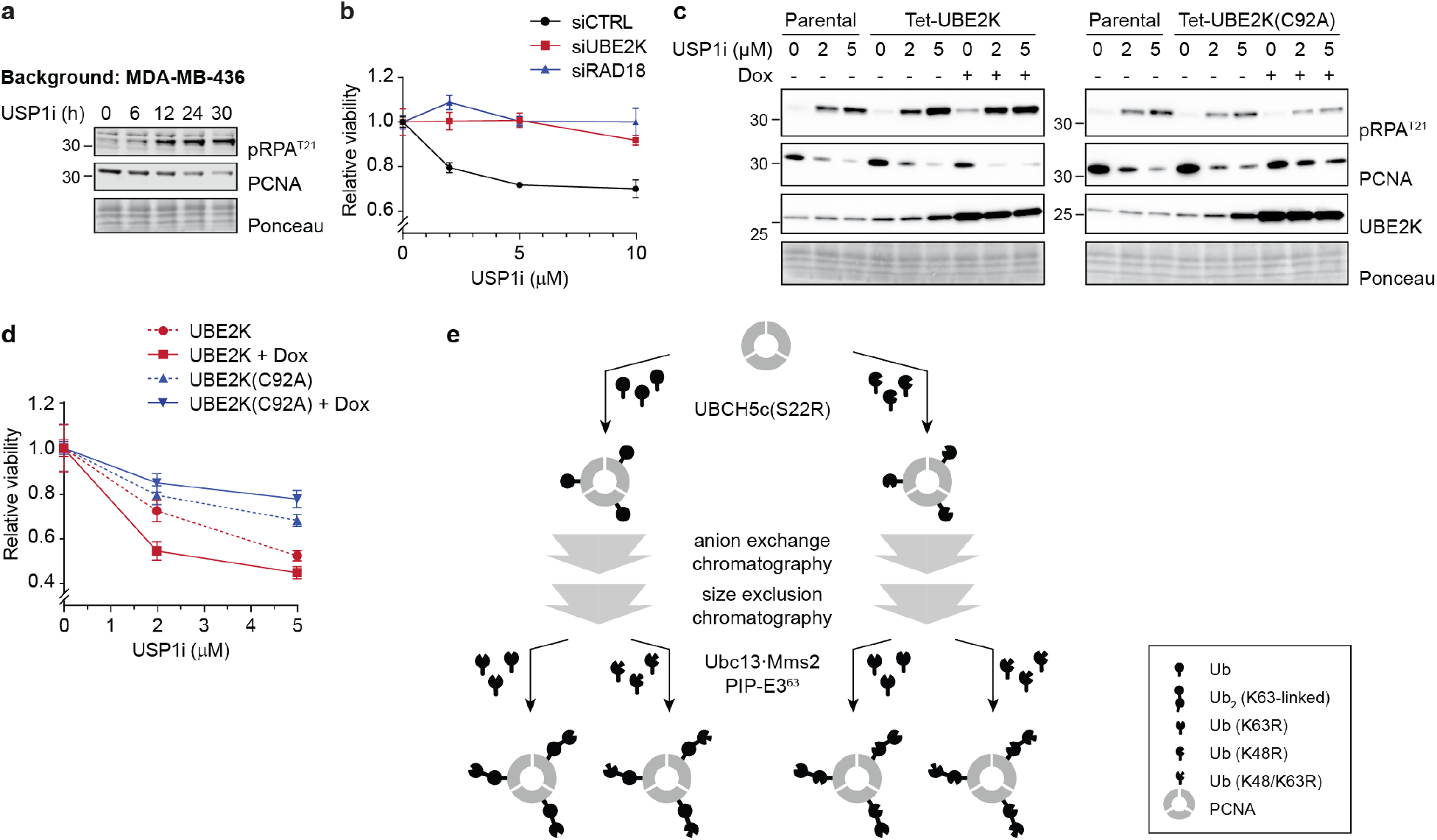
UBE2K mediates the DNA damage response in BRCA1-deficient cells. (**a**) USP1i causes damage signaling and PCNA degradation in BRCA1-deficient cells. MDA-MB-436 cells were treated with a USP1 inhibitor (30 µM ML-323) for the indicated periods, and checkpoint activation and PCNA were analyzed in whole cell lysates by western blotting. Ponceau S staining served as a loading control (pRPA^T21^: RPA32 phosphorylated at T21). (**b**) Depletion of UBE2K or RAD18 suppresses the toxic effect of USP1i. 48 h after siRNA transfection, MDA-MB-436 cells were treated with the indicated concentrations of ML-323 (USP1i) for 72 h, and viability was determined by the CellTiter-Blue Cell Viability Assay. Cell survival was normalized to the untreated condition for each knockdown. Data represent mean values and SD from three technical replicates. Shown is a representative assay of three independent experiments. (**c**) Overexpression of catalytically active UBE2K enhances damage signaling and PCNA degradation, while a catalytically inactive mutant (C92A) suppresses it. MDA-MB-436 cells harboring Dox-inducible UBE2K (WT or C92A) were treated simultaneously with 1 µg/ml Dox and the indicated concentrations of ML-323 (USPi) for 72 h. Western blot analysis was performed as in Fig. 1a. (**d**) Overexpression of a catalytically active UBE2K aggravates USP1i toxicity in BRCA1-deficient cells, while a catalytically inactve mutant (C92A) alleviates it. Viability was determined for the cells shown in panel **c** as described in panel **b**. (**e**) Schematic illustration of the ubiquitylation reactions and purification steps applied to prepare fully diubiquitylated PCNA carrying a K48R mutation at the proximal and/or distal positions for use in the reaction shown in Fig. 1c.

**Extended Data Fig. 2.**
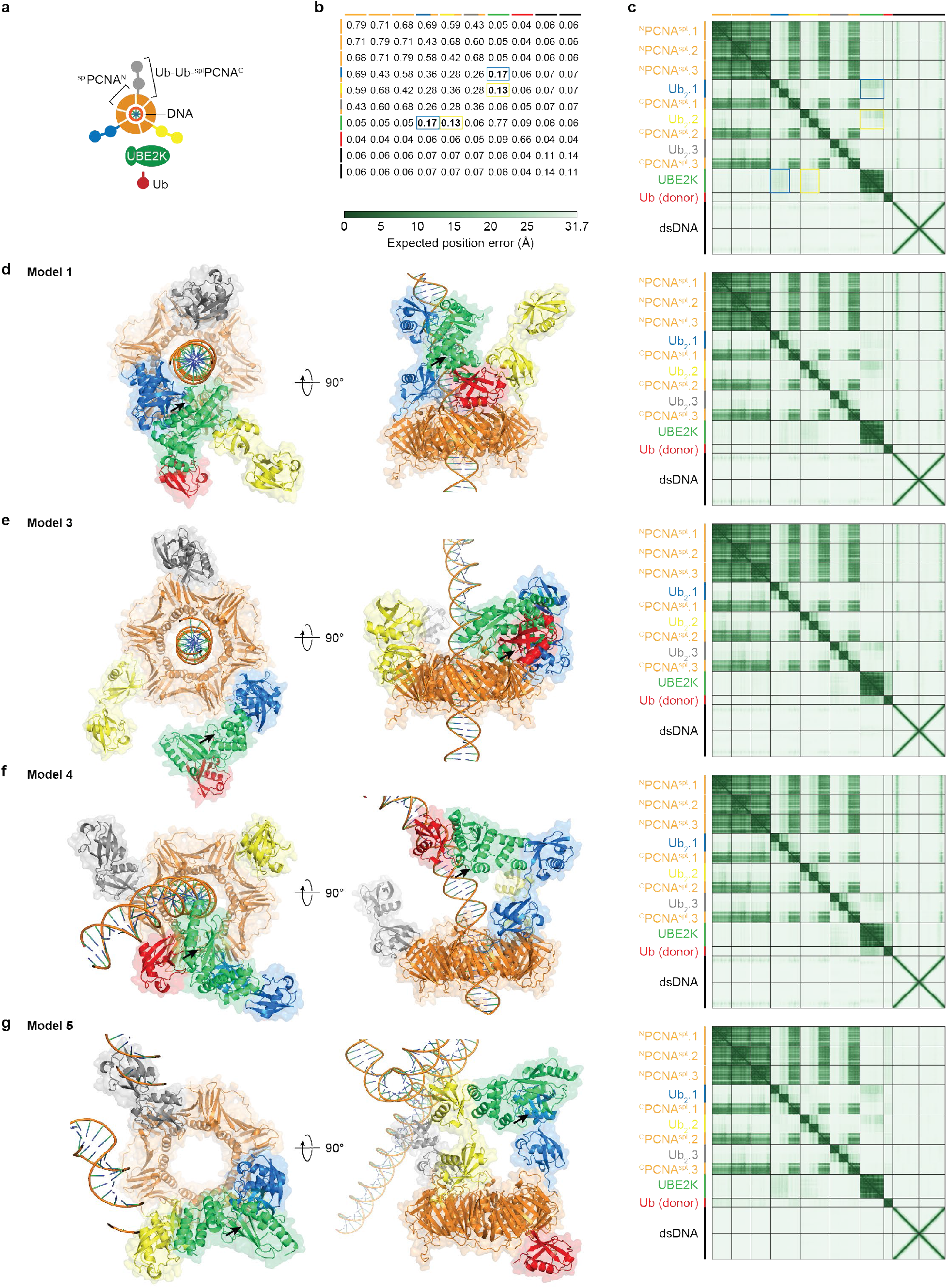
Trans-branching of K63-linked diubiquitin chains at the distal ubiquitin unit of DNA-loaded PCNA by UBE2K is sterically plausible. (**a**) A cartoon drawing illustrating the polypeptides used to mimic a complex of DNA-bound diubiquitylated PCNA with UBE2K and monoubiquitin as input for the AlphaFold 3 models shown here and in Fig. 1D (UBE2K-bound Ub_2_: blue; acceptor Ub_2_: yellow; donor Ub: red). ^spl^PCNA^N^ comprises amino acids 1-163 of PCNA; Ub-Ub-^spl^PCNA^C^ is a head-to-tail fusion of two ubiquitin moieties to amino acids 165-261 of PCNA (linked via Gly-Gly). Full sequences are provided in Extended Data Table 5. (**b**) Chain pair interface predicted template modelling (ipTM) score matrix of the AlphaFold 3 model shown in Fig. 1d. Blue and yellow boxes indicate potential interactions between UBE2K and two of the diubiquitin units. (**c**) Predicted aligned error (PAE) plot of the AlphaFold 3 model shown in Fig. 1d. Blue and yellow boxes indicate potential interactions between UBE2K and two of the diubiquitin units. (**d-g**) UBE2K is predicted to occupy various different positions on diubiquitylated PCNA. Top and side views as well as PAE plots of AlphaFold 3 models 1, 3, 4, and 5, generated along with model 2 using the same seed (Fig. 1d). Note that only part of the dsDNA is shown. In most models, UBE2K’s UBA domain contacts one of the distal ubiquitin units on PCNA. The consistent alignment of the ubiquitin chains parallel to the DNA axis on the back face of PCNA is compatible with a bridging of two chains by UBE2K. At the same time, UBE2K’s interactions with the ubiquitin chains do not obstruct the positioning of PCNA on DNA. In models 3 and 4 (as in model 2), the C-terminus of the free monoubiquitin (red) is juxtaposed to UBE2K’s catalytic cysteine, C92 (marked by black arrows), consistent with the role of a donor ubiquitin.

**Extended Data Fig. 3.**
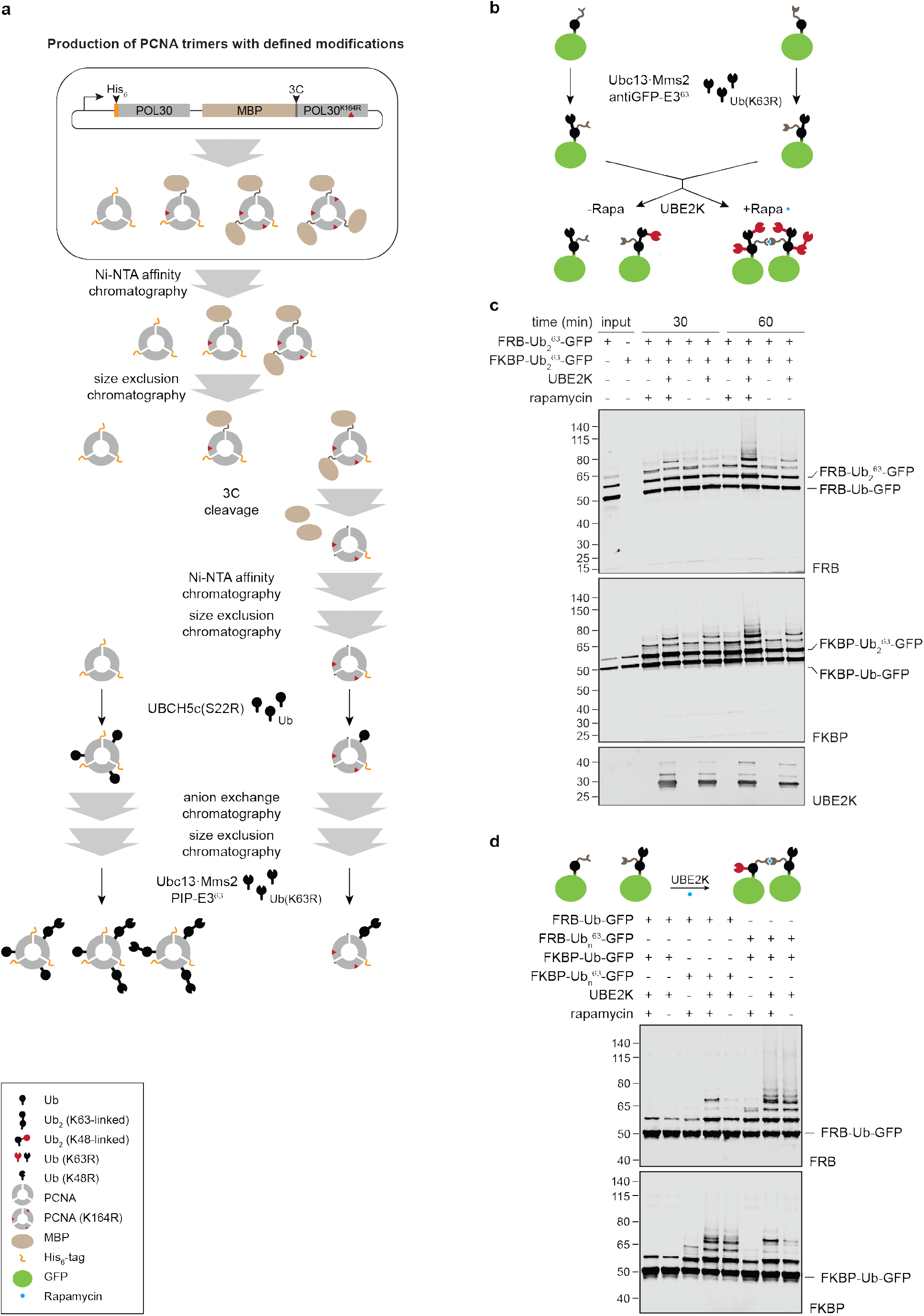
Purification schemes and *in vitro* reactions demonstrating trans-branching by UBE2K. (**a**) Schematic illustration of the production, purification, and ubiquitylation steps applied to prepare substrates for the *in vitro* reactions shown in Fig. 1e. Mixed trimers of PCNA were produced for a substrate carrying a single diubiquitin unit per trimer, and WT ^His^PCNA was produced separately to generate a substrate carrying up to three diubiquitin units. (**b**) Schematic illustration of the ubiquitylation reactions shown in Fig. 1f, used to demonstrate trans-activity of UBE2K on GFP model substrates capable of dimerizing via rapamycin-induced interactions between FRB and FKBP domains. (**c**) UBE2K preferentially branches K63-chains proximal to other chains. Branching was assessed *in vitro* according to the set-up shown in panel **b**, using K63-diubiquitylated Ub-GFP as substrates and inducing dimerization by rapamycin. The panel shows blots against the two substrates (FRB and FKBP) and UBE2K from the reactions shown in Fig. 1f. (**d**) UBE2K can extend monoubiquitin units proximal to a K63-chain. Chain extension was assessed *in vitro* using Ub-GFP and K63-diubiquitylated Ub-GFP substrate pairs fused to either FRB or FKBP, to allow for rapamycin-induced dimerization.

**Extended Data Fig. 4.**
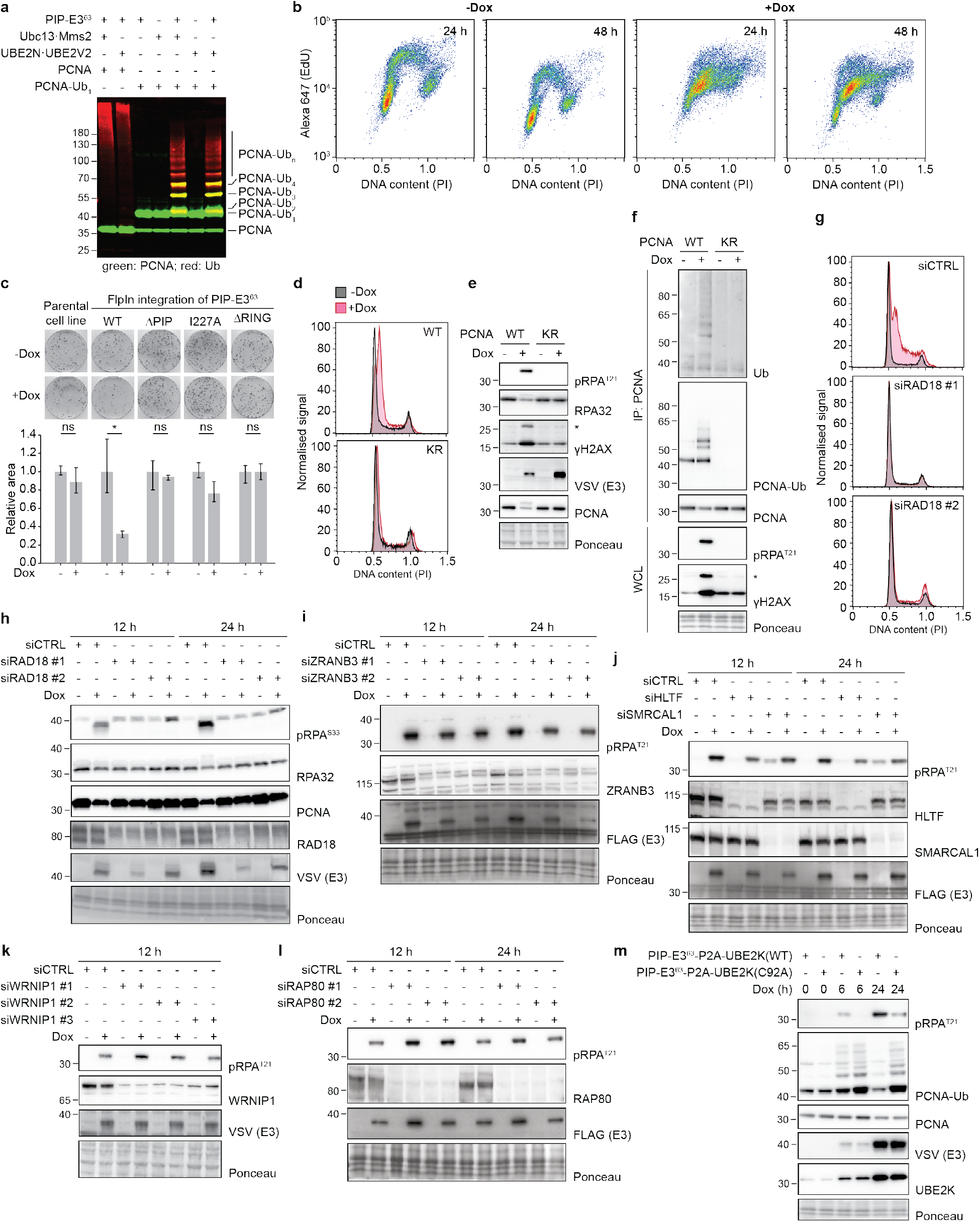
Unscheduled PCNA polyubiquitylation induces replication collapse and DNA damage signaling. (**a**) PIP-E3^63^ is active towards human PCNA monoubiquitylated at K164 with both yeast (Mms2·Ubc13) and human (UBE2V2·UBE2N) E2s. A blot of *in vitro* ubiquitylation reactions with recombinant proteins was probed against PCNA (green) and ubiquitin (red). (**b**) 2D replication profiles demonstrate replication collapse induced by PIP-E3^63^. RPE1 hTERT FlpIn PIP-E3^63^ cells were treated with 1 µg/ml doxycycline (Dox) for 24 h or 48 h. EdU (10 µM) was added 30 min prior to cell collection. DNA content and incorporated EdU were analyzed by flow cytometry (PI: propidium iodide). (**c**) Unscheduled PCNA polyubiquitylation compromises viability. Colony formation assays were performed on RPE1 hTERT FlpIn cells expressing PIP-E3^63^ or its mutant forms (+Dox). After 10 days of growth in the presence of 1 µg/ml Dox, representative images of fixed cells were taken, and the total area occupied by cells was quantified. For each cell line, the respective area was normalized to the uninduced (-Dox) condition. A representative set of two replicates is shown, indicating mean values and standard deviation. *: p=0.0261 (two-tailed t-test). (**d**) PIP-E3^63^-induced cell cycle arrest requires PCNA modification at K164. Cell cycle profiles were generated from wildtype (WT) and PCNA(K164R) mutant (KR) RPE1 hTERT cells harboring a doxycycline-inducible PIP-E3^63^ construct. (**e**) PIP-E3^63^-induced damage signaling requires PCNA modification at K164. Checkpoint activation was analyzed by western blotting of total lysates from the cells treated as shown in panel **d**. The asterisk shows the ubiquitylated form of ɣH2AX. (**f**) PCNA mono-and polyubiquitylation is absent in the KR cell line used in panels **d** and **e**. Ubiquitylated forms of PCNA were probed by western blotting of PCNA immunoprecipitates (IP). Damage markers were detected in whole cell lysates (WCL). The asterisk shows the ubiquitylated form of ɣH2AX. (**g**) PIP-E3^63^-induced S-phase arrest depends on RAD18. To monitor cell cycle profiles, RAD18 was transiently depleted in RPE1 hTERT FlpIn PIP-E3^63^ cells for 72 h by means of two independent siRNAs, followed by addition of 1 µg/ml Dox for 24 h. (**h**) PIP-E3^63^-induced damage signaling depends on RAD18. After depletion of RAD18 as in panel **g**, checkpoint activation, depletion efficiency, and E3 expression were analyzed after 12 and 24 h PIP-E3^63^ induction by western blotting of whole cell extracts. (**i**) ZRANB3 is not required for PIP-E3^63^-induced damage signaling. Checkpoint activation was analyzed after transiently depleting ZRANB3 analogous to the procedure shown in panel **h**. (**j**) HLTF and SMARCAL1 are not required for PIP-E3^63^-induced damage signaling. Checkpoint activation was analyzed analogous to the procedure shown in panel **h**. (**k**) WRNIP1 is not required for PIP-E3^63^-induced damage signaling. Checkpoint activation was analyzed analogous to the procedure shown in panel **h**. (**l**) RAP80 is not required for PIP-E3^63^-induced damage signaling. Checkpoint activation was analyzed analogous to the procedure shown in panel **h**. (**m**) Overexpression of catalytically inactive UBE2K(C92A) alleviates PIP-E3^63^-induced damage signaling and stabilizes ubiquitylated forms of PCNA. WT and mutant UBE2K were stably integrated in RPE1 hTERT cells as fusions to PIP-E3^63^ separated by a self-cleavable P2A peptide and expressed by addition of 1 µg/ml Dox for the indicated periods. Checkpoint markers, PCNA and enzymes were detected by western blotting of whole cell lysates.

**Extended Data Fig. 5.**
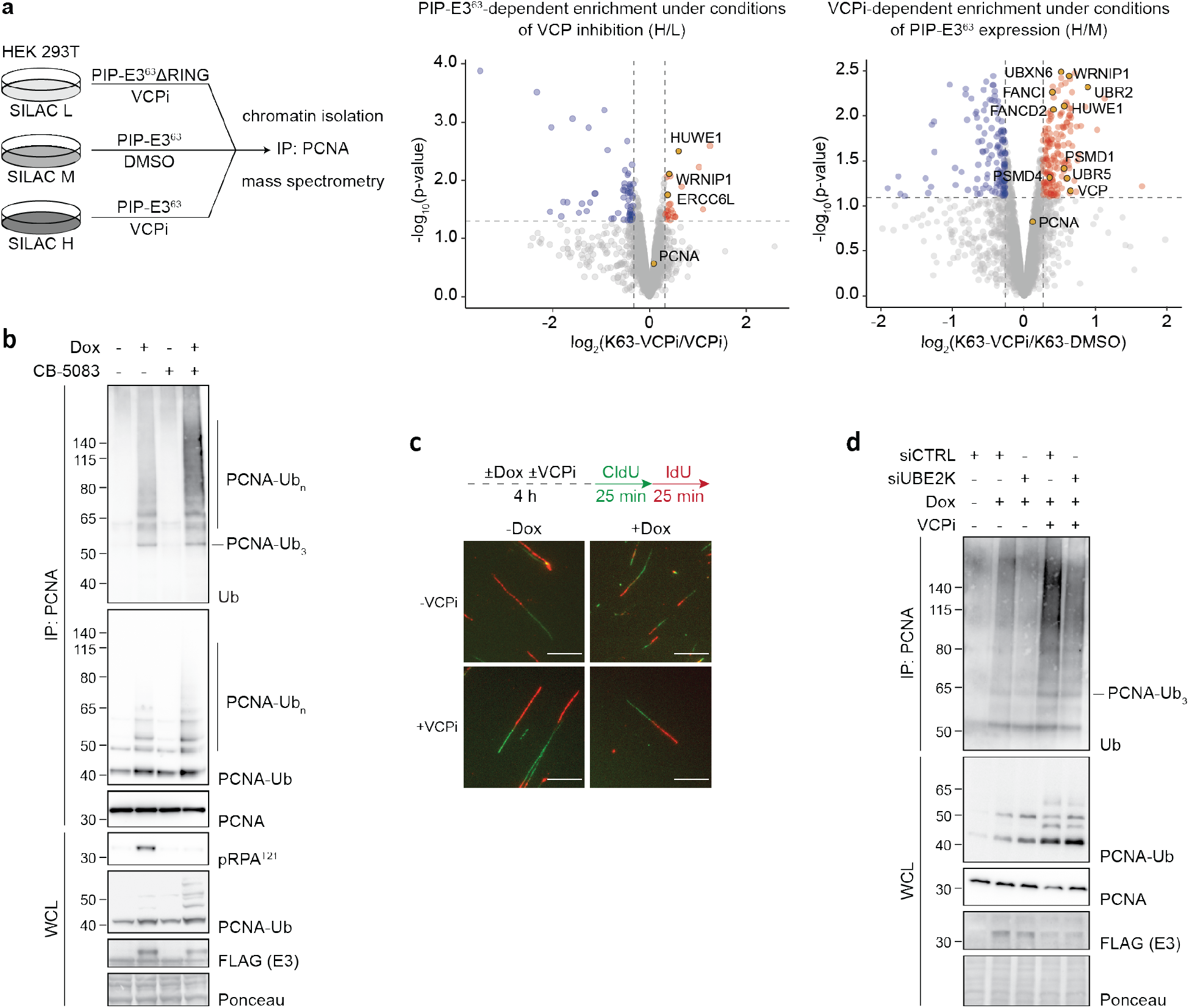
VCP mediates extraction of polyubiquitylated PCNA from chromatin. (**a**) SILAC-based mass spectrometry analysis identifies interactors of chromatin-bound PCNA polyubiquitylated by PIP-E3^63^ with or without a 6 h treatment with 5 μM NMS-873 (VCPi). The respective conditions are depicted on the scheme (left). Corresponding volcano plots for two pairwise comparisons are shown on the right. Selected interactors are highlighted and labeled. (**b**) An alternative VCP inhibitor, CB-5083, stabilizes polyubiquitylated PCNA on chromatin and prevents PIP-E3^63^-induced damage signaling. RPE1 hTERT FlpIn PIP-E3^63^ cells were treated with 1 µg/ml Dox and 0.5 μM CB-5083 or DMSO for 6 h. PCNA was immuno-precipitated from the chromatin fraction and its ubiquitylation was analyzed by western blotting. (**c**) Representative images of DNA fibers corresponding to Fig. 2I. Scale bar = 10 µm. (**d**) UBE2K promotes the formation of high-molecular weight ubiquitylated forms of PCNA. RPE1 hTERT FlpIn PIP-E3^63^ cells were transiently depleted of UBE2K for 72 h, followed by treatment with 1 µg/ml Dox and 5 µM NMS-873 (VCPi) as indicated for 6 h. PCNA was immunoprecipitated from the chromatin fraction and its ubiquitylation was analyzed by western blotting. Expression of the E3 and levels of unmodified and ubiquitylated PCNA were analyzed in total cell extracts.

**Extended Data Fig. 6.**
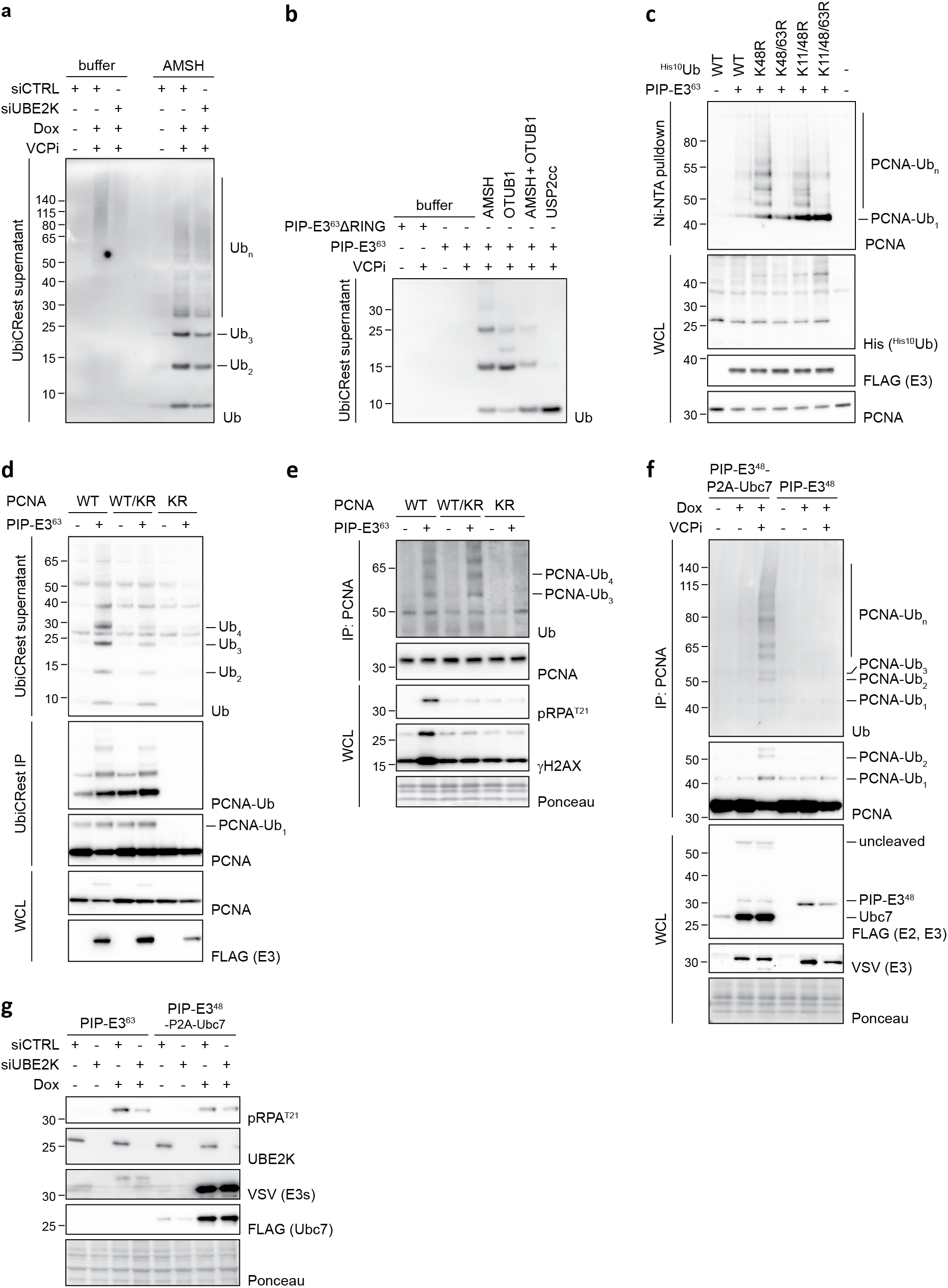
UBE2K promotes trans-branching on PCNA in cells. (**a**) Depletion of UBE2K reduces the amount of branched polyubiquitin chains on PCNA after induction of PIP-E3^63^. 72 h after siRNA transfection, RPE1 hTERT FlpIn PIP-E3^63^ cells were treated with 1 μg/ml Dox and 5 μM NMS-873 (VCPi) for 6 h. PCNA immunoprecipitated from the chromatin fraction was subjected to UbiCRest (treatment with 3 µM AMSH for 1 h) as in Fig. 3a,b. Cleaved ubiquitin species in the soluble fraction were analyzed by western blotting. (**b**) K48-linkages dominate among the branches on polyubiquitylated PCNA. UbiCRest assays were performed as in Fig. 3a,b. Alongside AMSH, the K48-selective DUB, OTUB1 (10 μM), and the catalytic domain of the linkage-insensitive DUB, USP2 (1 μM), were used. (**c**) Inhibition of K48-branching stabilizes polyubiquitylated PCNA. PIP-E3^63^ was transiently overexpressed in HEK 293T cells together with the indicated His_10_-tagged ubiquitin mutants. 48 h after transfection, tagged ubiquitin was pulled down from cells under denaturing conditions, followed by western blotting for PCNA. Expression of ^His^Ub and PIP-E3^63^ was analyzed by western blotting of total cell extracts. (**d**) Efficient branching requires multiple K63-chains on the PCNA trimer. PIP-E3^63^ was transiently overexpressed in HEK293T cells harboring different combinations of wild-type (WT) and K164R-mutant (KR) PCNA subunits. Following VCPi (5 µM NMS-873) for 3 h, UbiCRest was performed as in Fig. 3a,b. (**e**) PIP-E3^63^-mediated checkpoint activation requires ubiquitylation of multiple subunits of the PCNA trimer. HEK293T cells harboring different combinations of wild-type (WT) and mutant (KR) PCNA subunits with integrated PIP-E3^63^ were treated with 1 µg/ml Dox for 24 h. PCNA polyubiquitylation was analyzed by western blotting after PCNA precipitation from the chromatin fraction. Checkpoint activation and E3 expression were analyzed in whole cell lysates. (**f**) Co-expression of yeast Ubc7 boosts the activity of PIP-E3^48^ towards PCNA in human cells. RPE1 hTERT FlpIn cells harboring integrated PIP-E3^48^ alone or fused to Ubc7 via a self-cleavable peptide (PIP-E3^48^-P2A-Ubc7) were treated with 1 µg/ml Dox and VCPi or DMSO as indicated for 6 h. PCNA was immunoprecipitated from the chromatin fraction and its ubiquitylation was analyzed by western blotting. Expression of the enzymes was analyzed in whole cell extracts. Note that both PIP-E3^48^ and Ubc7 carry a FLAG-tag, while only the E3 carries a VSV-tag. (**g**) Residual checkpoint activation by direct K48-polyubiquitylation of PCNA is independent of UBE2K. RPE1 hTERT FlpIn PIP-E3^63^ and RPE1 hTERT FlpIn PIP-E3^48^-P2A-Ubc7 cells were transiently depleted of UBE2K for 72 h, followed by treatment with 1 µg/ml Dox for 24 h. Checkpoint activation (pRPA^T21^), depletion efficiency, and expression of the enzymes were analyzed in total cell extracts by western blotting.

**Extended Data Fig. 7.**
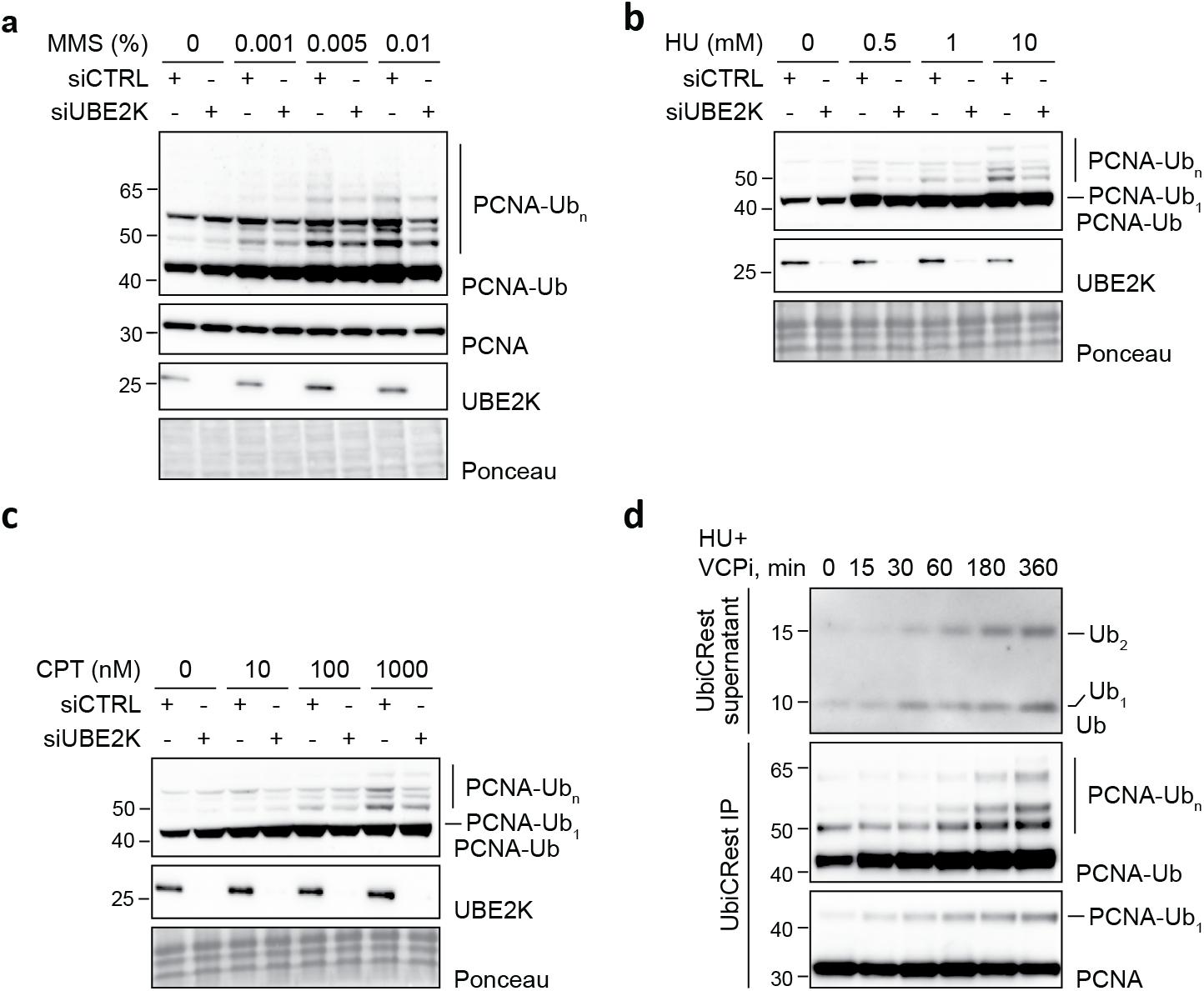
UBE2K modulates PCNA ubiquitylation in response to DNA replication stress. (**a**) UBE2K promotes PCNA polyubiquitylation in response to MMS. 72 h after siRNA transfection, RPE1 hTERT cells were treated with the indicated concentrations of MMS for 1 h, followed by release into medium containing 5 µM NMS-873 (VCPi) for 4h. PCNA ubiquitylation was analyzed in total cell extracts by western blotting. (**b**) UBE2K promotes PCNA polyubiquitylation in response to HU. 72 h after siRNA transfection, UBE2K RPE1 hTERT cells were treated with the indicated concentrations of HU for 24 h. PCNA ubiquitylation was analyzed in total cell extracts by western blotting. (**c**) UBE2K promotes PCNA polyubiquitylation in response to CPT. 72 h after siRNA transfection, RPE1 hTERT cells were treated with the indicated concentrations of CPT for 1 h, followed by release into medium containing 5 µM NMS-873 (VCPi) for 4 h. PCNA ubiquitylation was analyzed in total cell extracts by western blotting. (**d**) Branching on PCNA exhibits a time delay relative to K63-linked polyubiquitylation. RPE1 hTERT cells were treated with a combination of 4 mM HU and 5 µM NMS-873(VCPi) for the indicated periods, analogous to the assay shown in Fig. 4c, but in the absence of USP1i. PCNA was immunoprecipitated from the chromatin fraction and subjected to UbiCRest by treatment with 3 µM AMSH for 1 h, followed by analysis of supernatant and immobilized fractions (IP) by western blotting.

**Extended Data Fig. 8.**
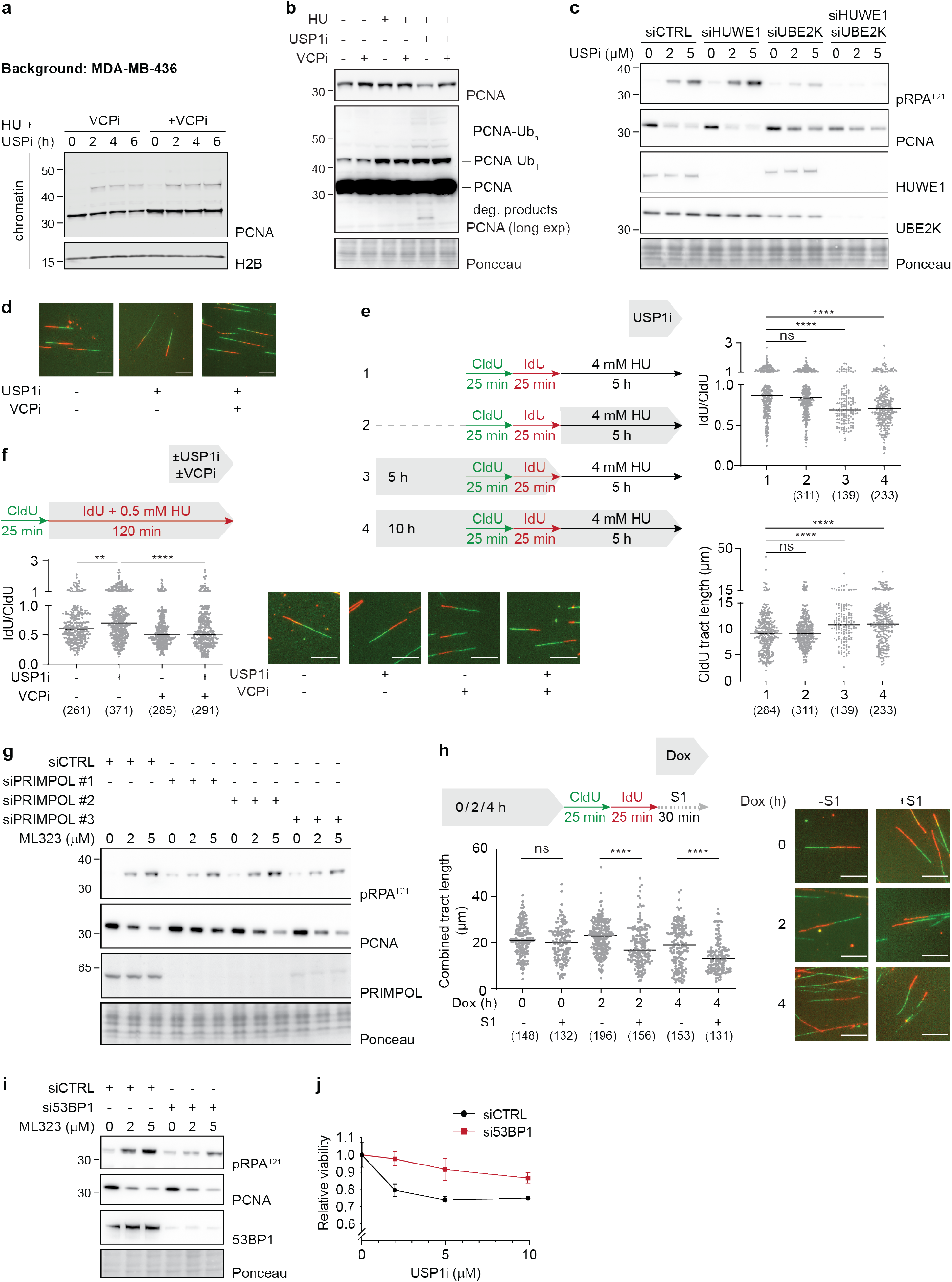
Synthetic lethality between BRCA1 and USP1 deficiency is caused by a hyper-accumulation of daughter-strand gaps. (**a**) USP1i-induced loss of chromatin-bound PCNA in BRCA1-deficient cells depends on VCP. MDA-MB-436 cells were treated with 4 mM HU, 30 µM ML-323 (USP1i) and 5 µM NMS-873 (VCPi) as indicated. PCNA levels were analyzed in the chromatin fraction by western blotting. Histone H2B served as loading control. (**b**) USP1i-induced degradation of PCNA in BRCA1-deficient cells depends on VCP. MDA-MB-436 cells were treated for 4 h with USP1i, VCPi, and 4 mM HU as indicated. Overall levels and ubiquitylation pattern of PCNA were analyzed by western blotting in total cell extracts. (**c**) Depletion of UBE2K but not HUWE1 prevents PCNA degradation and checkpoint activation in BRCA1-deficient cells. The indicated factors were depleted in MDA-MB-436 cells for 48 h, followed by treatment with USP1i as indicated for 72 h. Checkpoint activation (pRPA^T21^), total PCNA levels, and depletion efficiencies were analyzed by western blotting of total cell extracts. (**d**) Representative images of DNA fibers corresponding to the experiment described in Fig. 5e. Scale bar = 10 µm. (**e**) USP1i before but not during fork stalling results in nascent DNA degradation in BRCA1-deficient cells. MDA-MB-436 cells were treated as in Fig. 5d, but using 4 mM HU, followed by DNA spreading and fiber analysis. A representative experiment from two replicates is shown. Significance levels were calculated using the two-tailed Mann-Whitney test from the indicated number of fibers per sample (ns: p>0.05; ****: p<0.0001). (**f**) USP1i in BRCA1-deficient cells causes unrestrained fork progression upon replication stress in a VCP-dependent manner. MDA-MB-436 cells were treated as indicated, followed by DNA spreading and fiber analysis. Representative fiber images are shown. Scale bar = 10 µm. A representative experiment from three replicates is shown. Significance levels were calculated using the two-tailed Mann-Whitney test from the indicated number of fibers per sample (**: p = 0.0014; ****: p<0.0001). (**g**) Depletion of PRIMPOL has no impact on checkpoint activation and PCNA degradation in BRCA1-deficient cells. PRIMPOL was depleted in MDA-MB-436 cells for 48 h, followed by USP1i for 72 h. Checkpoint activation (pRPA^T21^), total PCNA levels, and depletion efficiency were analyzed by western blotting of total cell extracts. (**h**) PIP-E3^63^ expression causes an accumulation of daughter-strand gaps. RPE1 hTERT FlpIn PIP-E3^63^ cells were treated with 1 µg/ml Dox as indicated, followed by DNA fiber assays with S1 nuclease treatment. Representative fiber images are shown. Scale bar = 10 µm. Significance levels were calculated using the two-tailed Mann-Whitney test from the indicated number of fibers per sample (ns: p>0.05; ****: p<0.0001). (**i**) Depletion of 53BP1 reduces checkpoint activation without preventing PCNA degradation in BRCA1-deficient cells. 53BP1 was depleted in MDA-MB-436 cells for 48 h, followed by treatment with the indicated concentrations of ML-323 (USP1i) for 72 h. Checkpoint activation (pRPA^T21^), total PCNA levels, and depletion efficiency were analyzed by western blotting of total cell extracts. (**j**) Depletion of 53BP1 suppresses the toxic effect of USP1i. 48 h after siRNA transfection, MDA-MB-436 cells were treated with the indicated concentrations of ML-323 (USP1i) for 72 h, and viability was determined as in Extended Data Fig. 1b. Cell survival was normalized to the untreated condition for each knockdown. Data represent mean values and SD from three technical replicates. Shown is a representative assay of three independent experiments.

**Extended Data Table 1.**
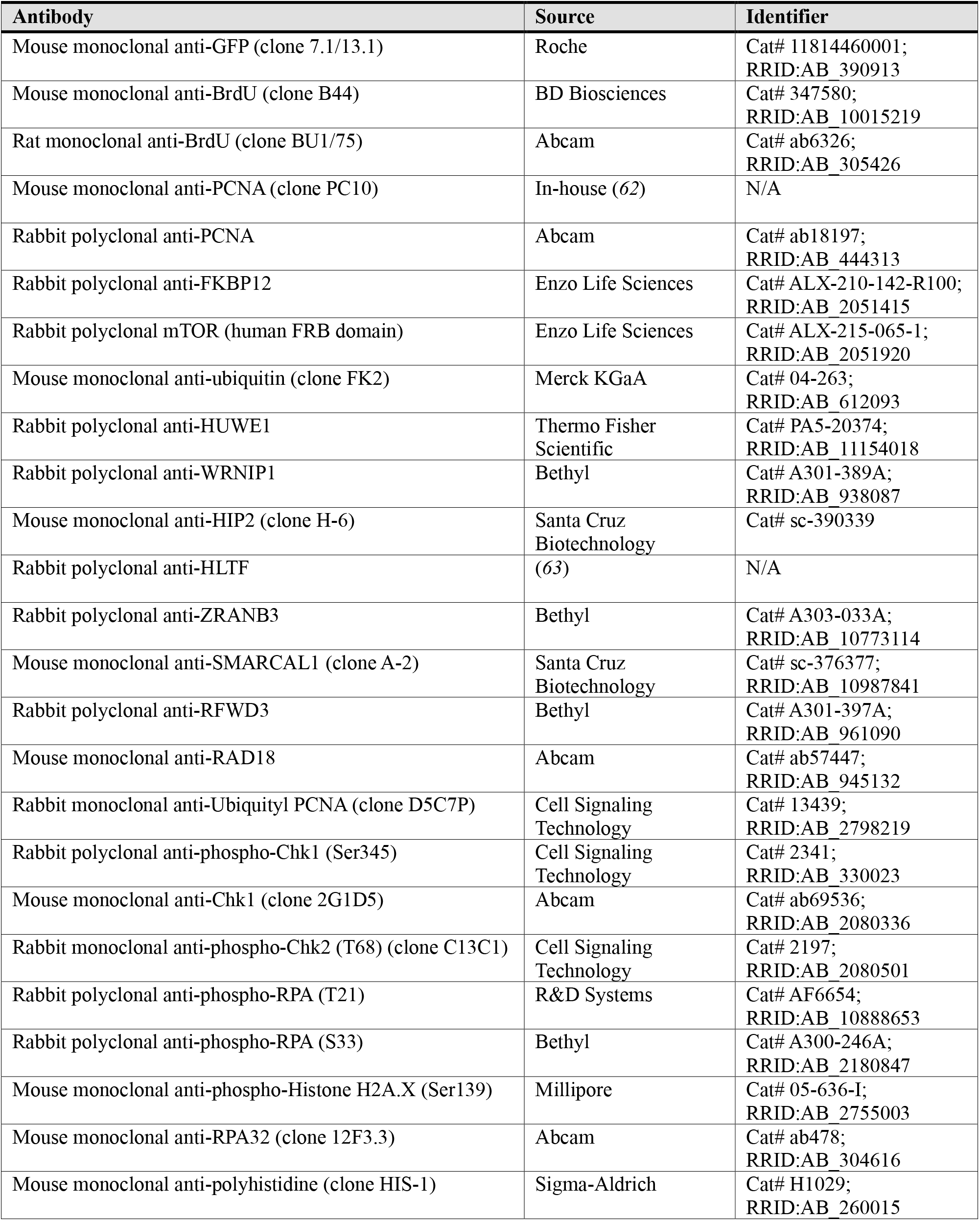
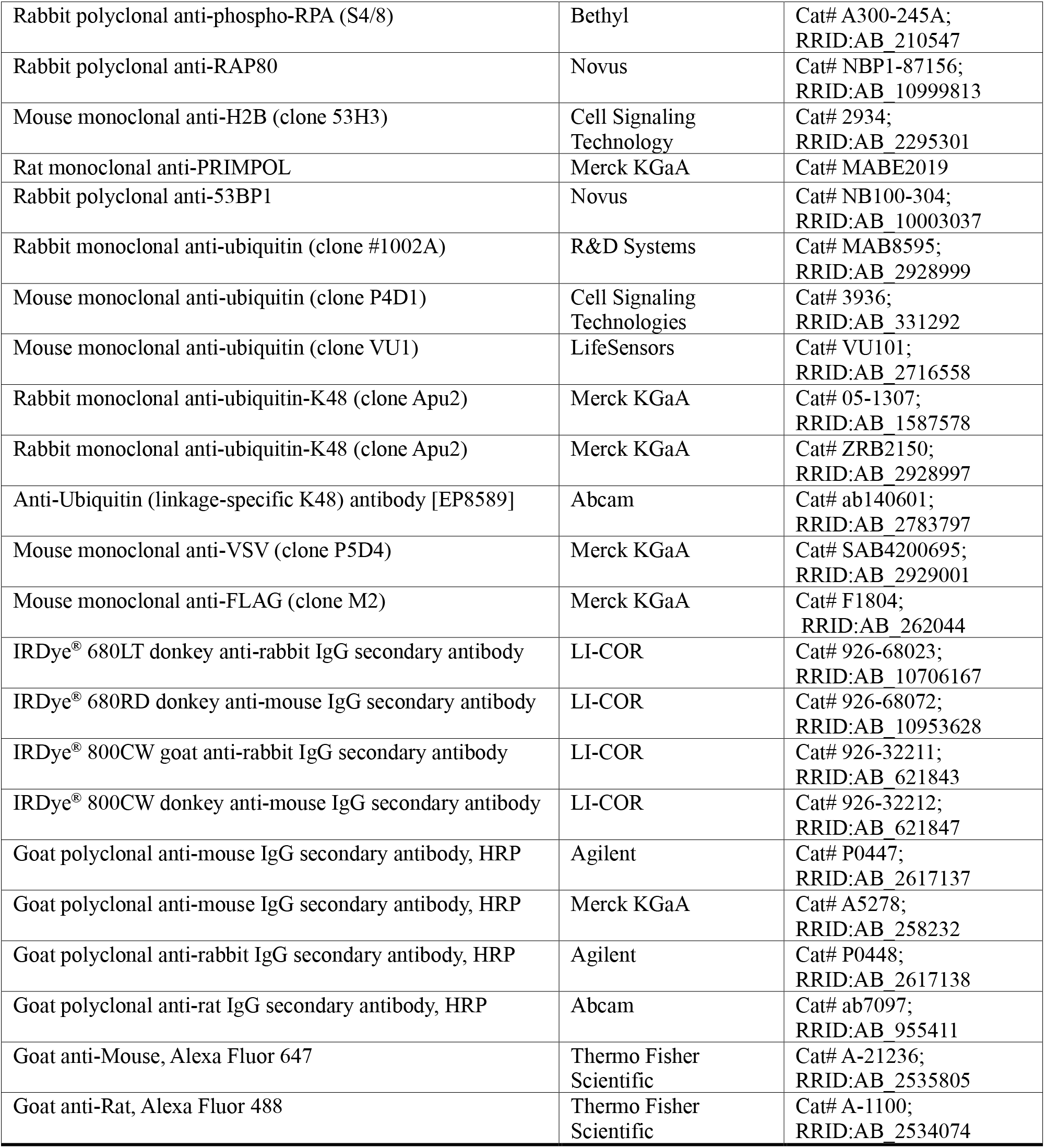
Antibodies used in this study.

**Extended Data Table 2.** Plasmids used in this study.

| Plasmid | ID | Use | Source |
| --- | --- | --- | --- |
| **Plasmids for use in mammalian cells** |  |  |  |
| pOG44 | 1809 | Creation of FlpIn cell lines | Gift from Simon Boulton<br>(Invitrogen Cat# V600520) |
| pMDLg/pRRE | 3805 | Production of lentivirus; packaging plasmid with Gag & Pol | Gift from Didier Trono (Addgene #12251) |
| pRSV-Rev | 3806 | Production of lentivirus; packaging plasmid with Rev | Gift from Didier Trono (Addgene #12253) |
| pMD2.G | 3807 | Production of lentivirus; VSV-G envelope expressing plasmid | Gift from Didier Trono (Addgene #12259) |
| pDEST/FRT/TO-FLAG-VSV-PIP-E3<sup>63</sup> | 4599 | Transient expression of <sup>FLAG-VSV</sup>PIP-E3<sup>63</sup>; creation of FlpIn cell lines | This study |
| pDEST/FRT/TO-FLAG-VSV-PIP-E3<sup>63</sup> (I227A) | 4715 | Transient expression of <sup>FLAG-VSV</sup>PIP-E3<sup>63</sup> (I227A); creation of FlpIn cell lines | This study |
| pDEST/FRT/TO-FLAG-VSV-E3<sup>63</sup> | 4720 | Transient expression of <sup>FLAG-VSV</sup>E3<sup>63</sup>; creation of FlpIn cell lines | This study |
| pDEST/FRT/TO-FLAG-VSV-PIP-E3<sup>63</sup>ΔRING | 4756 | Transient expression of <sup>FLAG-VSV</sup>PIP-E3<sup>63</sup>ΔRING; creation of FlpIn cell lines | This study |
| pLentiCMV rtTA3 Blast | 4892 | Constitutive expression of the reverse tetracycline transactivator rtTA3 | Gift from Eric Campeau (Addgene #26429) |
| pLentiCMV/Tre/3G Neo-VSV-PIP-E3<sup>63</sup> | 4894 | Creation of stable cell lines with inducible expression of <sup>VSV</sup>PIP-E3<sup>63</sup> | This study |
| pDEST/FRT/TO-FLAG-VSV-PIP-E3<sup>48</sup>-FLAG | 5112 | Transient expression of <sup>FLAG-VSV</sup>PIP-E3<sup>48</sup>-Flag; creation of FlpIn cell lines | This study |
| pDEST/FRT/TO-FLAG-VSV-PIP-E3<sup>48</sup>-P2A-3xFLAG-hcoUbc7 | 5136 | Transient expression of <sup>FLAG-VSV</sup>PIP-E3<sup>48</sup>-P2A-<sup>3xFLAG</sup>Ubc7; creation of FlpIn cell lines | This study |
| pDEST-His10-Ub | 5242 | Transient expression of <sup>His</sup>Ub | This study |
| pDEST-His10-Ub(K48R) | 5340 | Transient expression of <sup>His</sup>Ub<sup>K48R</sup> | This study |
| pDEST-His10-Ub(K63R) | 5341 | Transient expression of <sup>His</sup>Ub<sup>K63R</sup> | This study |
| pDEST-His10-Ub(K48/63R) | 5342 | Transient expression of <sup>His</sup>Ub<sup>K48/63R</sup> | This study |
| pDEST-His10-Ub(K11/48/63R) | 5346 | Transient expression of <sup>His</sup>Ub<sup>K11/48/63R</sup> | This study |
| pDEST-His10-Ub(K11/48R) | 5347 | Transient expression of <sup>His</sup>Ub<sup>K11/48R</sup> | This study |
| pLentiCMV/Tre/3G Neo-PIP-E3<sup>63</sup>-P2A-UBE2K | 6390 | Creation of stable cell lines with inducible expression of PIP-E3<sup>63</sup>-P2A-UBE2K | This study |
| pLentiCMV/Tre/3G Neo-PIP-E3<sup>63</sup>-P2A-UBE2K(C92A) | 6394 | Creation of stable cell lines with inducible expression of PIP-E3<sup>63</sup>-P2A-UBE2K(C92A) | This study |
| pLentiCMV/Tre/3G Neo-UBE2K | 6422 | Creation of stable cell lines with inducible expression of UBE2K | This study |
| pLentiCMV/Tre/3G Neo-UBE2K(C92A) | 6423 | Creation of stable cell lines with inducible expression of UBE2K(C92A) | This study |
| **Plasmids for production of recombinant proteins** |  |  |  |
| pGEX-6P-1-AMSH | 2898 | Production of AMSH | (33) |
| pOPINK-OTUB1 | 3456 | Production of OTUB1 | (33) |
| pET15b-His6-TEV-USP2cc | 3239 | Production of <sup>His</sup>USP2cc | (57) |
| pRSF-duet-His6-PCNA | 2095 | Production of human <sup>His</sup>PCNA | Gift from Svend Petersen-Mahrt (this study) |
| pRSF-duet-His6-PCNA-MBP-3C-hPCNA(K164R) | 6323 | Production of mixed PCNA trimers | This study |
| pET3a-Ub(K63R) | 684 | Production of Ub<sup>K63R</sup> | (64) |
| pET28-His6-mE1 | 2448 | Production of <sup>His</sup>Uba1 (mouse) | (54) |
| pGEX-4T1-PIP-E3(63) | 2452 | Production of <sup>GST</sup>PIP-E3<sup>63</sup> | (28) |
| pET30-His6-TEV-Mms2 | 2807 | Production of Mms2 (*S. cerevisiae*) | (56) |
| pET21(+)-His6-Ubc13 | 3010 | Production of <sup>His</sup>Ubc13 (*S. cerevisiae*) | (56) |
| pGEX-4T1-Ubc13 | 0347 | Production of GSTUbc13 (*S. cerevisiae*) | (59) |
| pET30-His6-TEV-UBE2K | 3357 | Production of <sup>His</sup>UBE2K | This study |
| pET30-His6-TEV-UBCH5c(S22R) | 6106 | Production of UBCH5c(S22R) | (58) |
| pET24a-antiGFP-L20-E3(63)-His6 | 6290 | Production of antiGFP-E3<sup>63</sup>-His | This study |
| pET15-His6-FKBP-L20-3C-Ub(G76V)-GFP | 5409 | Production of HisFKBP-Ub-GFP | This study |
| pET15-His6-FRB(L2098T)-L20-3C-Ub(G76V)-GFP | 6362 | Production of HisFRB-Ub-GFP | This study |

**Extended Data Table 3.**
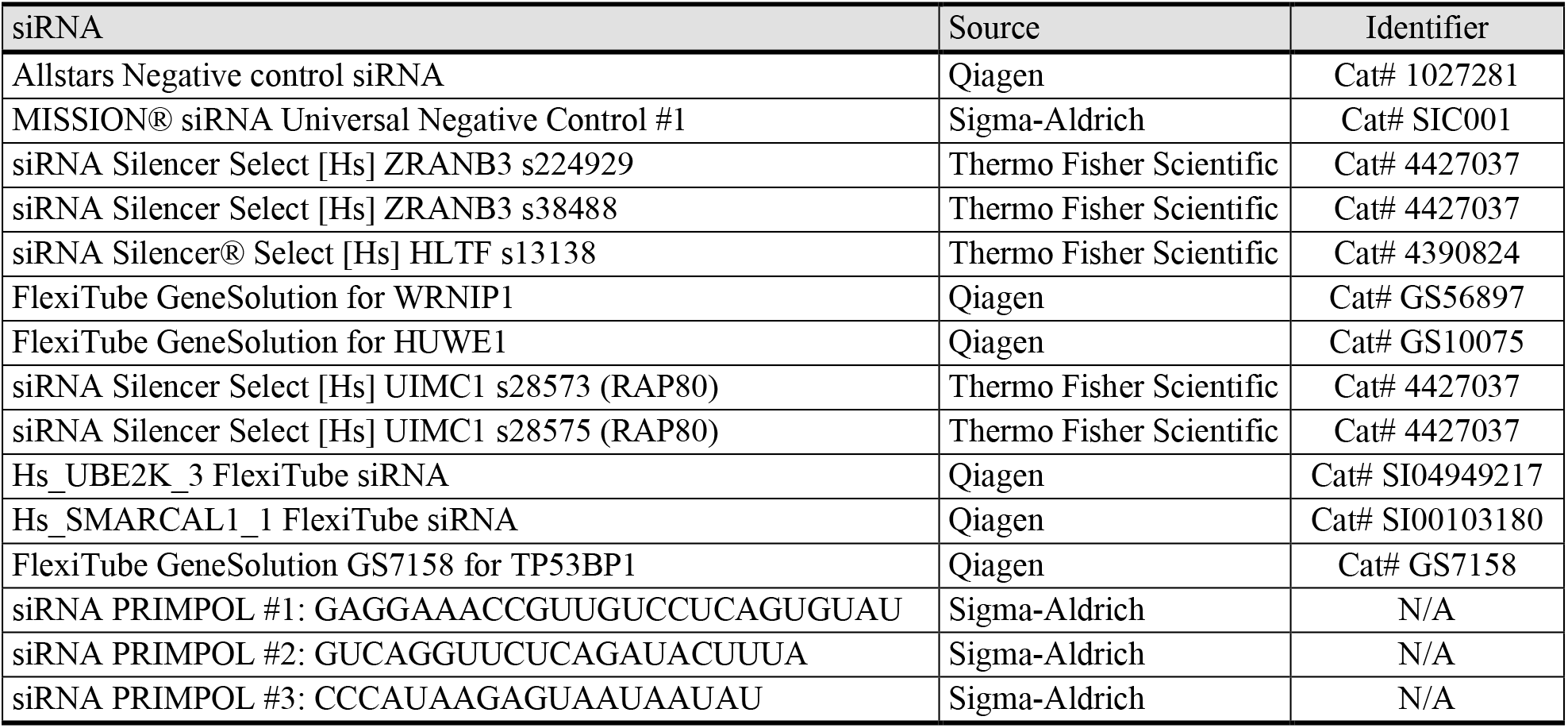
siRNAs used in this study.

**Extended Data Table 4.** Bacterial strains and cell lines used in this study.

| Strain or cell line | Source | Identifier |
| --- | --- | --- |
| **Bacterial strain** |  |  |
| *Escherichia coli* Top10 | Thermo Fisher Scientific | Cat# C404010 |
| *Escherichia coli* BL21(DE3) | Merck KGaA | Cat# 69450-3 |
| *Escherichia coli* BL21(DE3)pLysS | Merck KGaA | Cat# 69451-3 |
| *Escherichia coli* BL21-CodonPlus (DE3)-RIL | Agilent | Cat# 230245 |
| *Escherichia coli* Rosetta™ 2(DE3)pLysS | Merck KGaA | Cat# 71403-3 |
| *Escherichia coli* LOBSTR-BL21(DE3)-RIL | Kerafast | Cat# EC1002 |
| **Cell line** |  |  |
| MDA-MB-436 | ATCC | Cat# HTB-130 |
| MDA-MB-436 TetOn UBE2K | This study | N/A |
| MDA-MB-436 TetOn UBE2K(C92A) | This study | N/A |
| HEK 293T | Merck KGaA | Cat# 12022001-1VL<br>RRID:CVCL_0063 |
| HEK 293T PIP-E3(63) | This study | N/A |
| HEK 293T PCNA WT/K164R | (29) | N/A |
| HEK 293T PCNA WT/K164R PIP-E3(63) | This study | N/A |
| HEK 293T PCNA K164R | (29) | N/A |
| HEK 293T PCNA K164R PIP-E3(63) | This study | N/A |
| RPE1 hTERT WT | (29) | N/A |
| RPE1 hTERT WT PIP-E3(63) | This study | N/A |
| RPE1 hTERT PCNA K164R clone 2B10 | (29) | N/A |
| RPE1 hTERT PCNA K164R clone 2B10 PIP-E3(63) | This study | N/A |
| RPE1 hTERT WT PIP-E3(63)-P2A-UBE2K | This study | N/A |
| RPE1 hTERT WT PIP-E3(63)-P2A-UBE2K(C92A) | This study | N/A |
| RPE1 hTERT WT FRT | (65) | N/A |
| RPE1 hTERT WT FRT PIP-E3(63) | This study | N/A |
| RPE1 hTERT WT FRT PIP-E3(63) I227A | This study | N/A |
| RPE1 hTERT WT FRT PIP-E3(63)ΔRING | This study | N/A |
| RPE1 hTERT WT FRT E3(63) | This study | N/A |
| RPE1 hTERT WT FRT PIP-E3(48) | This study | N/A |
| RPE1 hTERT WT FRT PIP-E3(48)-P2A-Ubc7 | This study | N/A |
| RPE1 hTERT WT FRT PIP-E3(48)-P2A-Ubc7 | This study | N/A |

**Extended Data Table 5.**
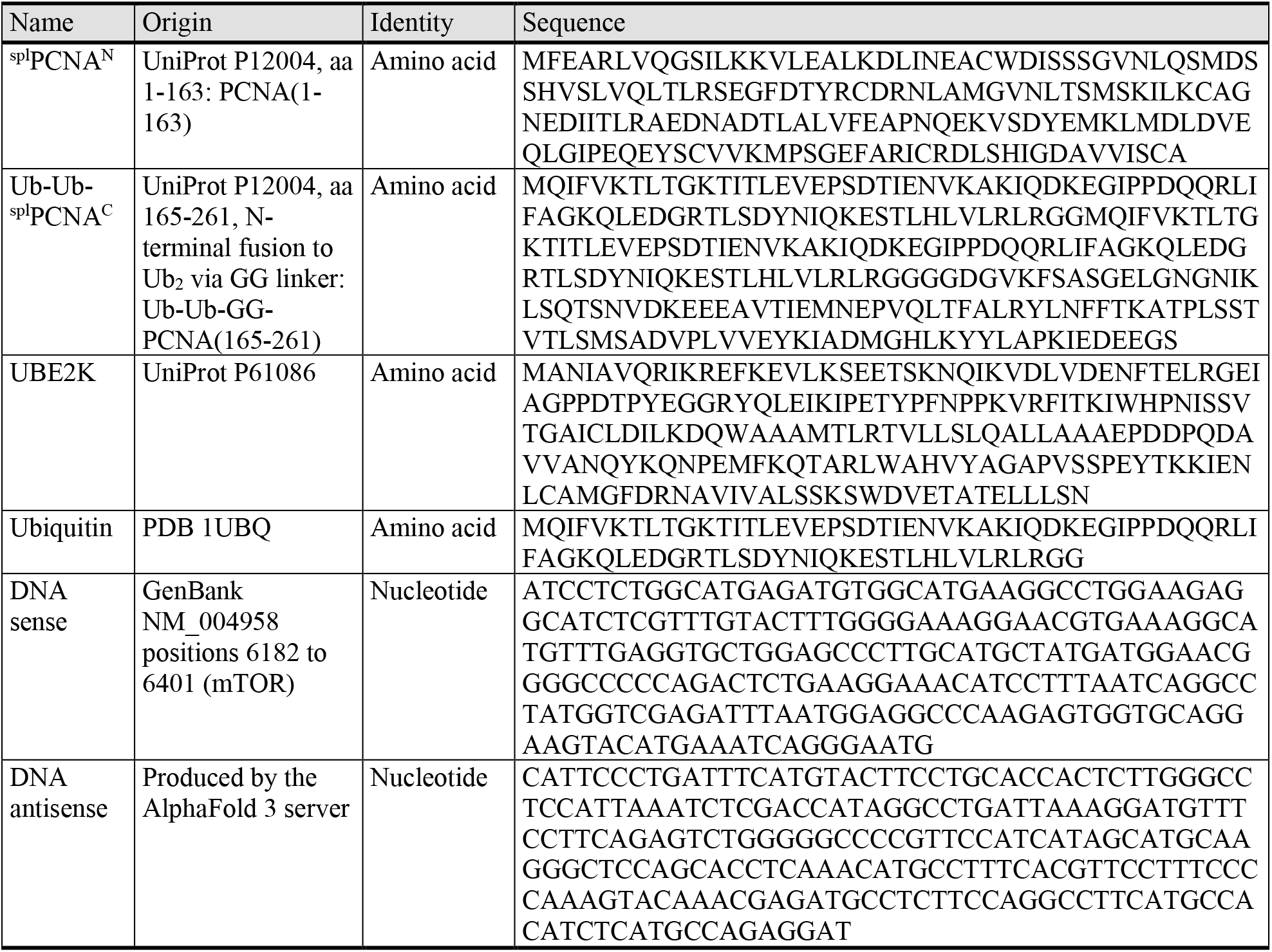
AlphaFold 3 input sequences used in this study.

## References

1. D. Komander, M. Rape, The ubiquitin code. Annu Rev Biochem 81, 203–229 (2012). doi:10.1146/annurev-biochem-060310-170328

2. Y. T. Kwon, A. Ciechanover, The Ubiquitin Code in the Ubiquitin-Proteasome System and Autophagy. Trends Biochem Sci 42, 873–886 (2017). doi:10.1016/j.tibs.2017.09.002

3. S. O. Crowe, A. Rana, K. K. Deol, Y. Ge, E. R. Strieter, Ubiquitin Chain Enrichment Middle-Down Mass Spectrometry Enables Characterization of Branched Ubiquitin Chains in Cellulo. Anal Chem 89, 4428–4434 (2017). doi:10.1021/acs.analchem.6b03675

4. F. Ohtake, Branched ubiquitin code: from basic biology to targeted protein degradation. J Biochem 171, 361–366 (2022). doi:10.1093/jb/mvac002

5. K. N. Swatek et al., Insights into ubiquitin chain architecture using Ub-clipping. Nature 572, 533–537 (2019). doi:10.1038/s41586-019-1482-y

6. E. E. Blythe, K. C. Olson, V. Chau, R. J. Deshaies, Ubiquitin-and ATP-dependent unfoldase activity of P97/VCP*NPLOC4*UFD1L is enhanced by a mutation that causes multisystem proteinopathy. Proc Natl Acad Sci U S A 114, E4380–E4388 (2017). doi:10.1073/pnas.1706205114

7. A. J. Boughton, S. Krueger, D. Fushman, Branching via K11 and K48 Bestows Ubiquitin Chains with a Unique Interdomain Interface and Enhanced Affinity for Proteasomal Subunit Rpn1. Structure 28, 29–43 e26 (2020). doi:10.1016/j.str.2019.10.008

8. S. M. Lange et al., VCP/p97-associated proteins are binders and debranching enzymes of K48-K63-branched ubiquitin chains. Nat Struct Mol Biol 31, 1872–1887 (2024). doi:10.1038/s41594-024-01354-y

9. H. J. Meyer, M. Rape, Enhanced protein degradation by branched ubiquitin chains. Cell 157, 910–921 (2014). doi:10.1016/j.cell.2014.03.037

10. F. Ohtake, Y. Saeki, S. Ishido, J. Kanno, K. Tanaka, The K48-K63 Branched Ubiquitin Chain Regulates NF-kappaB Signaling. Mol Cell 64, 251–266 (2016). doi:10.1016/j.molcel.2016.09.014

11. I. E. Wertz et al., Phosphorylation and linear ubiquitin direct A20 inhibition of inflammation. Nature 528, 370–375 (2015). doi:10.1038/nature16165

12. C. H. Emmerich et al., Activation of the canonical IKK complex by K63/M1-linked hybrid ubiquitin chains. Proc Natl Acad Sci U S A 110, 15247–15252 (2013). doi:10.1073/pnas.1314715110

13. J. van den Boom, H. Meyer, VCP/p97-Mediated Unfolding as a Principle in Protein Homeostasis and Signaling. Mol Cell 69, 182–194 (2018). doi:10.1016/j.molcel.2017.10.028

14. H. Tsuchiya et al., In Vivo Ubiquitin Linkage-type Analysis Reveals that the Cdc48-Rad23/Dsk2 Axis Contributes to K48-Linked Chain Specificity of the Proteasome. Mol Cell 66, 488–502 e487 (2017). doi:10.1016/j.molcel.2017.04.024

15. S. P. Jackson, D. Durocher, Regulation of DNA damage responses by ubiquitin and SUMO. Mol Cell 49, 795–807 (2013). doi:10.1016/j.molcel.2013.01.017

16. C. Hoege, B. Pfander, G. L. Moldovan, G. Pyrowolakis, S. Jentsch, RAD6-dependent DNA repair is linked to modification of PCNA by ubiquitin and SUMO. Nature 419, 135–141 (2002). doi:10.1038/nature00991

17. M. Bienko et al., Ubiquitin-binding domains in Y-family polymerases regulate translesion synthesis. Science 310, 1821–1824 (2005). doi:10.1126/science.1120615

18. P. Stelter, H. D. Ulrich, Control of spontaneous and damage-induced mutagenesis by SUMO and ubiquitin conjugation. Nature 425, 188–191 (2003). doi:10.1038/nature01965

19. A. Ciccia et al., Polyubiquitinated PCNA recruits the ZRANB3 translocase to maintain genomic integrity after replication stress. Mol Cell 47, 396–409 (2012). doi:10.1016/j.molcel.2012.05.024

20. M. Vujanovic et al., Replication Fork Slowing and Reversal upon DNA Damage Require PCNA Polyubiquitination and ZRANB3 DNA Translocase Activity. Mol Cell 67, 882–890 e885 (2017). doi:10.1016/j.molcel.2017.08.010

21. K. S. Lim et al., USP1 Is Required for Replication Fork Protection in BRCA1-Deficient Tumors. Mol Cell 72, 925–941 e924 (2018). doi:10.1016/j.molcel.2018.10.045

22. A. Simoneau et al., Ubiquitinated PCNA Drives USP1 Synthetic Lethality in Cancer. Mol Cancer Ther 22, 215–226 (2023). doi:10.1158/1535-7163.MCT-22-0409

23. A. da Costa et al., Single-stranded DNA Gap Accumulation is a Functional Biomarker for USP1 Inhibitor Sensitivity. Cancer Res 84, 3435–3446 (2024). doi:10.1158/0008-5472.CAN-23-4007

24. I. Garcia-Santisteban, G. J. Peters, E. Giovannetti, J. A. Rodriguez, USP1 deubiquitinase: cellular functions, regulatory mechanisms and emerging potential as target in cancer therapy. Mol Cancer 12, 91 (2013). doi:10.1186/1476-4598-12-91

25. C. E. Moore et al., RFWD3 promotes ZRANB3 recruitment to regulate the remodeling of stalled replication forks. J Cell Biol 222, (2023). doi:10.1083/jcb.202106022

26. L. Pluska et al., The UBA domain of conjugating enzyme Ubc1/Ube2K facilitates assembly of K48/K63-branched ubiquitin chains. EMBO J 40, e106094 (2021). doi:10.15252/embj.2020106094

27. J. Abramson et al., Accurate structure prediction of biomolecular interactions with AlphaFold 3. Nature 630, 493–500 (2024). doi:10.1038/s41586-024-07487-w

28. S. Wegmann et al., Linkage reprogramming by tailor-made E3s reveals polyubiquitin chain requirements in DNA-damage bypass. Mol Cell 82, 1589–1602 e1585 (2022). doi:10.1016/j.molcel.2022.02.016

29. T. Thakar et al., Ubiquitinated-PCNA protects replication forks from DNA2-mediated degradation by regulating Okazaki fragment maturation and chromatin assembly. Nat Commun 11, 2147 (2020). doi:10.1038/s41467-020-16096-w

30. L. A. Poole, D. Cortez, Functions of SMARCAL1, ZRANB3, and HLTF in maintaining genome stability. Crit Rev Biochem Mol Biol 52, 696–714 (2017). doi:10.1080/10409238.2017.1380597

31. N. Kanu et al., RAD18, WRNIP1 and ATMIN promote ATM signalling in response to replication stress. Oncogene 35, 4009–4019 (2016). doi:10.1038/onc.2015.427

32. B. Sobhian et al., RAP80 targets BRCA1 to specific ubiquitin structures at DNA damage sites. Science 316, 1198–1202 (2007). doi:10.1126/science.1139516

33. M. K. Hospenthal, T. E. T. Mevissen, D. Komander, Deubiquitinase-based analysis of ubiquitin chain architecture using Ubiquitin Chain Restriction (UbiCRest). Nat Protoc 10, 349–361 (2015). doi:10.1038/nprot.2015.018

34. J. McCullough, M. J. Clague, S. Urbe, AMSH is an endosome-associated ubiquitin isopeptidase. J Cell Biol 166, 487–492 (2004). doi:10.1083/jcb.200401141

35. T. Wang et al., Evidence for bidentate substrate binding as the basis for the K48 linkage specificity of otubain 1. J Mol Biol 386, 1011–1023 (2009). doi:10.1016/j.jmb.2008.12.085

36. A. M. Catanzariti, T. A. Soboleva, D. A. Jans, P. G. Board, R. T. Baker, An efficient system for high-level expression and easy purification of authentic recombinant proteins. Protein Sci 13, 1331–1339 (2004). doi:10.1110/ps.04618904

37. C. C. Chen, W. Feng, P. X. Lim, E. M. Kass, M. Jasin, Homology-Directed Repair and the Role of BRCA1, BRCA2, and Related Proteins in Genome Integrity and Cancer. Annu Rev Cancer Biol 2, 313–336 (2018). doi:10.1146/annurev-cancerbio-030617-050502

38. K. Cong, S. B. Cantor, Exploiting replication gaps for cancer therapy. Mol Cell 82, 2363–2369 (2022). doi:10.1016/j.molcel.2022.04.023

39. G. Nagaraju, R. Scully, Minding the gap: the underground functions of BRCA1 and BRCA2 at stalled replication forks. DNA Repair (Amst) 6, 1018–1031 (2007). doi:10.1016/j.dnarep.2007.02.020

40. G. Bai et al., HLTF Promotes Fork Reversal, Limiting Replication Stress Resistance and Preventing Multiple Mechanisms of Unrestrained DNA Synthesis. Mol Cell 78, 1237–1251 e1237 (2020). doi:10.1016/j.molcel.2020.04.031

41. D. J. Martins, S. Tirman, A. Quinet, C. F. M. Menck, Detection of Post-Replicative Gaps Accumulation and Repair in Human Cells Using the DNA Fiber Assay. J Vis Exp, (2022). doi:10.3791/63448

42. L. J. Bainbridge, R. Teague, A. J. Doherty, Repriming DNA synthesis: an intrinsic restart pathway that maintains efficient genome replication. Nucleic Acids Res 49, 4831–4847 (2021). doi:10.1093/nar/gkab176

43. K. Cong et al., Replication gaps are a key determinant of PARP inhibitor synthetic lethality with BRCA deficiency. Mol Cell 81, 3128–3144 e3127 (2021). doi:10.1016/j.molcel.2021.06.011

44. E. Cybulla et al., A RAD18-UBC13-PALB2-RNF168 axis mediates replication fork recovery in BRCA1-deficient cancer cells. Nucleic Acids Res 52, 8861–8879 (2024). doi:10.1093/nar/gkae563

45. M. A. Nakasone et al., Structure of UBE2K-Ub/E3/polyUb reveals mechanisms of K48-linked Ub chain extension. Nat Chem Biol 18, 422–431 (2022). doi:10.1038/s41589-021-00952-x

46. A. Waltho et al., K48-and K63-linked ubiquitin chain interactome reveals branch-and length-specific ubiquitin interactors. Life Sci Alliance 7, e202402740 (2024). doi:10.26508/lsa.202402740

47. C. Renz et al., Ubiquiton-An inducible, linkage-specific polyubiquitylation tool. Mol Cell 84, 386–400 e311 (2024). doi:10.1016/j.molcel.2023.11.016

48. A. Kirchhofer et al., Modulation of protein properties in living cells using nanobodies. Nat Struct Mol Biol 17, 133–138 (2010). doi:10.1038/nsmb.1727

49. A. Shevchenko, H. Tomas, J. Havlis, J. V. Olsen, M. Mann, In-gel digestion for mass spectrometric characterization of proteins and proteomes. Nat Protoc 1, 2856–2860 (2006). doi:10.1038/nprot.2006.468

50. J. Cox, M. Mann, MaxQuant enables high peptide identification rates, individualized p.p.b.-range mass accuracies and proteome-wide protein quantification. Nat Biotechnol 26, 1367–1372 (2008). doi:10.1038/nbt.1511

51. J. Cox et al., Andromeda: a peptide search engine integrated into the MaxQuant environment. J Proteome Res 10, 1794–1805 (2011). doi:10.1021/pr101065j

52. J. E. Elias, S. P. Gygi, Target-decoy search strategy for increased confidence in large-scale protein identifications by mass spectrometry. Nat Methods 4, 207–214 (2007). doi:10.1038/nmeth1019

53. M. E. Ritchie et al., limma powers differential expression analyses for RNA-sequencing and microarray studies. Nucleic Acids Res 43, e47 (2015). doi:10.1093/nar/gkv007

54. A. F. Carvalho et al., High-yield expression in Escherichia coli and purification of mouse ubiquitin-activating enzyme E1. Mol Biotechnol 51, 254–261 (2012). doi:10.1007/s12033-011-9463-x

55. R. G. Hibbert, T. K. Sixma, Intrinsic flexibility of ubiquitin on proliferating cell nuclear antigen (PCNA) in translesion synthesis. J Biol Chem 287, 39216–39223 (2012). doi:10.1074/jbc.M112.389890

56. C. Renz et al., Ubc13-Mms2 cooperates with a family of RING E3 proteins in budding yeast membrane protein sorting. J Cell Sci 133, (2020). doi:10.1242/jcs.244566

57. J. Shi et al., Nuclear myosin VI maintains replication fork stability. Nat Commun 14, 3787 (2023). doi:10.1038/s41467-023-39517-y

58. L. P. Swift et al., SNM1A is crucial for efficient repair of complex DNA breaks in human cells. Nat Commun 15, 5392 (2024). doi:10.1038/s41467-024-49583-5

59. H. D. Ulrich, Protein-protein interactions within an E2-RING finger complex. Implications for ubiquitin-dependent DNA damage repair. J Biol Chem 278, 7051–7058 (2003). doi:10.1074/jbc.M212195200

60. B. D. Freudenthal, L. Gakhar, S. Ramaswamy, M. T. Washington, Structure of monoubiquitinated PCNA and implications for translesion synthesis and DNA polymerase exchange. Nat Struct Mol Biol 17, 479–484 (2010). doi:10.1038/nsmb.1776

61. C. Elfmann, J. Stülke, PAE viewer: a webserver for the interactive visualization of the predicted aligned error for multimer structure predictions and crosslinks. Nucleic Acids Res 51, W404–W410 (2023). doi:10.1093/nar/gkad350

62. N. H. Waseem, D. P. Lane, Monoclonal antibody analysis of the proliferating cell nuclear antigen (PCNA). Structural conservation and the detection of a nucleolar form. J Cell Sci 96 (Pt 1), 121–129 (1990). doi:10.1242/jcs.96.1.121

63. G. Debauve et al., Early expression of the Helicase-Like Transcription Factor (HLTF/SMARCA3) in an experimental model of estrogen-induced renal carcinogenesis. Mol Cancer 5, 23 (2006). doi:10.1186/1476-4598-5-23

64. J. L. Parker, H. D. Ulrich, Mechanistic analysis of PCNA poly-ubiquitylation by the ubiquitin protein ligases Rad18 and Rad5. EMBO J 28, 3657–3666 (2009). doi:10.1038/emboj.2009.303

65. J. Mansfeld, P. Collin, M. O. Collins, J. S. Choudhary, J. Pines, APC15 drives the turnover of MCC-CDC20 to make the spindle assembly checkpoint responsive to kinetochore attachment. Nat Cell Biol 13, 1234–1243 (2011). doi:10.1038/ncb2347

